# Mechanism of how the universal module XMAP215 γ-TuRC nucleates microtubules

**DOI:** 10.1101/2024.06.03.597159

**Authors:** Collin T. McManus, Sophie M. Travis, Philip D. Jeffrey, Rui Zhang, Sabine Petry

**Author notes:** Correspondence and requests for materials should be addressed to Sabine Petry.

## Abstract

It has become increasingly evident in recent years that nucleation of microtubules from a diverse set of MTOCs requires both the γ-tubulin ring complex (γ-TuRC) and the microtubule polymerase XMAP215. Despite their essentiality, little is known about how these nucleation factors interact and work together to generate microtubules. Using biochemical domain analysis of XMAP215 and structural approaches, we find that a sixth TOG domain in XMAP215 binds γ-TuRC *via* γ-tubulin as part of a broader interaction involving the C-terminal region. Moreover, TOG6 is required for XMAP215 to promote nucleation from γ-TuRC to its full extent. Interestingly, we find that XMAP215 also depends strongly on TOG5 for microtubule lattice binding and nucleation. Accordingly, we report a cryo-EM structure of TOG5 bound to the microtubule lattice that reveals promotion of lateral interactions between tubulin dimers. Finally, we find that while XMAP215 constructs’ effects on nucleation are generally proportional to their effects on polymerization, formation of a direct complex with γ-TuRC allows cooperative nucleation activity. Thus, we propose that XMAP215’s C-terminal TOGs 5 and 6 play key roles in promoting nucleation by promoting formation of longitudinal and lateral bonds in γ-TuRC templated nascent microtubules at cellular MTOCs.

## INTRODUCTION

Microtubules are hollow, cylindrical protein polymers assembled from ɑβ-tubulin heterodimers. They are involved in a range of essential cellular functions including intracellular organization and transport, resilience to mechanical stress, cell motility, and chromosome segregation. Although purified tubulin will form, or nucleate, microtubules spontaneously at sufficient concentrations in vitro, cells regulate where and when microtubule nucleation occurs through localization of nucleation factors^1,2^. These factors aid nucleation by promoting assembly of tubulin oligomers. Essential for the nucleation of microtubules in a eukaryotic cell is the microtubule nucleation module consisting of the γ-tubulin ring complex (γ-TuRC) and *Xenopus* microtubule assembly protein of 215kDa (XMAP215) / ch-TOG ^3–6^.

γ-TuRC promotes nucleation by templating tubulin assembly in a geometry roughly matching the helical parameters of a microtubule ^7–11^. It presents a ring of 14 γ-tubulin monomers, each supported by one of five γ-tubulin complex proteins (GCPs 2-6). Importantly, γ-TuRC’s overall architecture is splayed open compared to the microtubule such that the template is imperfect with low activity in vitro ^10,12,13^. Nevertheless, computational modeling, nucleation assays using purified components, and structural studies have demonstrated that assembly of tubulin onto the γ-TuRC template provides sufficient lateral bond energy to close the γ-TuRC ring, a conformation in which spoke 14 overlaps and occludes spoke 1^10,12,14,15^. γ-TuRC thus consistently nucleates and caps 13 protofilament microtubules ^14–16^. γ-TuRC was thought to be the only essential nucleator until it was shown that XMAP215 is equally important, and moreover acts synergistically with γ-TuRC, an activity that had been previously posited two decades ago^4,17,18^.

XMAP215 is a plus-end tracking protein that promotes microtubule polymerization by recruiting soluble tubulin ^19,20^. Tubulin binding by XMAP215 orthologs is achieved by a variable number of tumor overexpressed gene (TOG) domains; arrays of two TOG domains have been identified in yeast, three in *C. elegans*, and five in other eukaryotes like *D. melanogaster* and vertebrates^21^. Crystal structures from yeast, *C. elegans*, *D. melanogaster*, and human orthologs provide a full complement of structural models of the five XMAP215 TOG domains ^22–29^. These domains share similar architectures, containing 12 helices in six HEAT repeats (HRs) A-F that stack to form a 60 Å alpha-solenoid paddle. Conserved residues in the intra-HEAT repeat loops mediate binding to tubulin on the surface that is exposed on the microtubule exterior ^23–25,29,30^. Despite their homology, the TOG domains in pentameric XMAP215 orthologs have structural differences conserved at their positions in the TOG array that are thought to establish XMAP215’s orientation on the microtubule plus end and allow for processive polymerization^31^. Polymerization activity has been further linked to microtubule lattice affinity conferred by the flexible linkers following TOG2 and TOG4^30,32^.

More recently, it was found that XMAP215 is also required for microtubule nucleation, that XMAP215 directly binds to γ-TuRC, and that both components form a module that synergistically nucleates a microtubule in the cell ^4,5,33,34^. Although co-recruitment may often be transient, XMAP215 has also been reported to colocalize with γ-TuRC *in vitro* and at centriolar subdistal appendages, where it may play a role in anchoring microtubules ^4,5,12^. This anchoring activity is consistent with XMAP215’s ability to nucleate microtubule asters when tethered to resin via C-terminal tags ^18,35^. However, precisely how XMAP215 contributes to microtubule nucleation with γ-TuRC and generates the MT cytoskeleton of the cell is yet to be uncovered.

In this study, we characterize the structures and activities of multiple domains to reveal how XMAP215 helps nucleate microtubules with γ-TuRC. We show that there is indeed a TOG6 domain C-terminal to TOG5 in XMAP215, and that, while TOG6 plays a role that specifically enhances the efficiency of nucleation, TOG5 functions to promote both nucleation and polymerization. Our work uncovers general rules of how polymerization and nucleation are linked and helps explain how the microtubule cytoskeleton is formed.

## RESULTS

### XMAP215’s C-terminal region contains a TOG domain with unique architecture

XMAP215 and γ-TuRC interact via XMAP215’s C-terminal region in a manner yet to be determined. Based on secondary structure prediction and solution NMR data of the *D. melanogaster* XMAP215 ortholog,^36,37^ we hypothesized that a sixth TOG domain (TOG6) resides within XMAP215’s C-terminal domain, and that it is critical for binding specifically to γ-tubulin rather than ɑβ-tubulin like its N-terminal structural homologs.

First, we used AlphaFold2^38^ to predict the structure of the *Xenopus laevis* full length XMAP215 (Fig. S1a). Consistent with prior structural data, the prediction includes five canonical TOG domains, each connected by flexible linkers. Strikingly, downstream of TOG5 in what has previously been referred to as the XMAP215 C-terminal domain, an additional domain was predicted. It is also composed of 6 pairs of alpha helices arranged in a paddle, consistent with general TOG morphology, and thus provides further support for the putative TOG6 domain (Fig. 1a, S1a-b), though it is the most divergent from the other XMAP215 TOG domains (Fig. S1a, d). To assess this prediction, we crystalized TOG6 and collected a moderate resolution dataset (∼6Å) which we solved by molecular replacement (Data not shown). This revealed density consistent with the helices in the TOG6 prediction (Fig. 1b). In addition, a four-helix bundle stands out against the unstructured linkers and individual alpha-helices of the rest of the C-terminal domain prediction (Fig. 1a, S1c). This domain, which we termed “extended” or abbreviated “ext”, occurs 18 amino acids downstream of TOG6 and resembles the last 4 helices of some TOG domains (Fig. S1c).

**Fig. 1.**
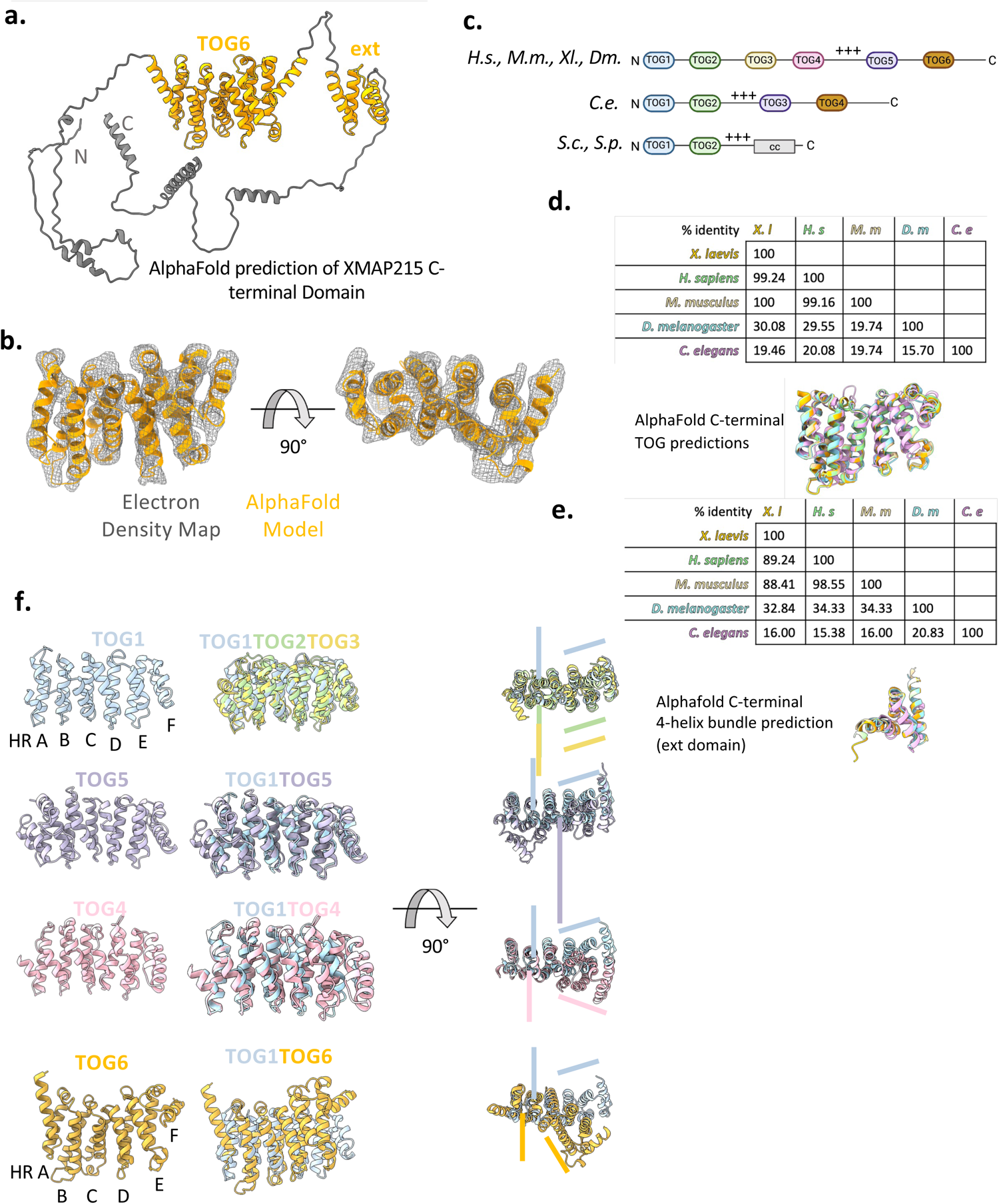
A unique TOG6 domain exists in the C-terminal region of XMAP215. **a)** AlphaFold2 prediction of the XMAP215 C-terminal region including all residues after TOG5 (aa1463-2065). TOG6 and the four helix bundle “ext” domain are shown in orange. Other residues are gray. **b)** TOG6 AlphaFold2 prediction superimposed on the electron density map derived by X-ray crystallography using molecular replacement. Resolution about 5.7Å. **c)** Schematics of XMAP215 orthologs from *Homo sapiens (H.s.), Mus musculus (M.m), Xenopus laevis (X.l.), Drosophila melanogaster (D.m), Caenorhabditis elegans (C.e.), Saccharomyces cerevisiae (S.c.), and Saccharomyces pombe (S.p.)* “+++” indicates positively changed linker domains. Created with Biorender.com **d)** Percent identity matrix from Clustal Omega alignment of TOG6 homologous domains from the higher order eukaryotic orthologs in C and superimposition of those domains^56^. **e)** Percent identity matrix from Clustal Omega alignment of four helix bundle “ext” homologous domains from the higher order eukaryotic orthologs in C and superimposition of those domains^56^. **f)** TOG domains 1, 2, 3, 4, 5, and 6 extracted from the AlphaFold2 prediction for full length *X. laevis* XMAP215 and superimposed for comparison. N-termini are positioned to the left, and C-termini to the right. In the images to the left of the arrow, the canonical tubulin binding surfaces are positioned downwards. In the rotated images to the right of the arrow, the non-tubulin binding surface is shown. Colored lines illustrate the kink or lack thereof which occurs between heat repeats C and D.

To assess whether the TOG6 domain is evolutionarily conserved, we performed AlphaFold2 predictions for XMAP215 orthologs across species. We found TOG6 domains within the C-terminal domains of flies, humans, and mice (Fig. 1c-d). Remarkably, an analogous TOG4 in *C. elegans* ZYG-9, which has only three previously described TOG domains and whose third TOG domain corresponds to XMAP215 TOG5^25,28^, as well as the less conserved fly Msps TOG6, also resulted in the same predicted fold (Fig. 1c-d). This suggests that this domain has a meaningful and specific function requiring its structural conservation on par with the other TOG domains. Moreover, the extended four helix bundle downstream of TOG6 is conserved as well across these species (Fig. 1e), highlighting its functional importance yet to be determined.

Given that XMAP215’s TOG domains have unique structures that are conserved with respect to their positions in the TOG array, we asked how TOG6 compares to XMAP215’s other TOG domains. For comparison, we aligned the six TOG domains from the *X. laevis* XMAP215 AlphaFold2 prediction (Fig. S1a) on the best conserved heat repeats A-C (Fig. 1f) (Byrnes et al., 2017). The most extreme difference between TOG6 and the other TOG domains is in the kink in the alpha solenoid stack that occurs between heat repeats C and D (Fig. 1f). When viewed from the non-tubulin binding surface, TOGs1, 2, and 3 exhibit similar kinks, with planes generated from heat repeats A-C and D-F intersecting at −28° (Fig. S1d). TOG4 exhibits a kink in the opposite direction but of similar magnitude with an angle of +23°. The kink in TOG5 of −7.6° is negligible, giving it an overall straight geometry as previously noted (Byrnes et al., 2017). Surprisingly the kink in TOG6 greatly exaggerated at +69° (Fig. S1d). We also noticed that TOG6 heat repeats A, B, and C appear more staggered than in the other TOG domains, with a shorter heat repeat A sitting higher, and the longer heat repeats B and C reaching lower (Fig. 1f). In sum, the unique structures of TOG domains 4, 5 and 6 imply they may have unique functions.

### TOG6 specifically binds γ-tubulin as part of a broader interaction between XMAP215’s C-terminal region and γ-TuRC

Having defined the structure of TOG6, we next tested whether TOG6 is involved in XMAP215’s interaction with γ-TuRC. We designed several GFP fusions of XMAP215 truncation and deletion constructs (Fig. 2a) and assessed their binding to γ-tubulin in a pull-down assay. A construct containing TOG5 through the C-terminus (TOG5-C) but not TOG5 was able to bind to γ-tubulin, indicating that the binding region resides in the C-terminal domain (Fig. 2b). TOG6 binds γ-tubulin to a similar extent as TOG6 through the four helix bundle (TOG6ext) and the C-terminal domain (C-term), which contains all residues downstream of TOG5 (Fig. 2b). A TOG6ext deletion from the C-terminal domain (CΔTOG6ext) exhibits decreased binding to γ-tubulin, suggesting that this region is important for binding γ-tubulin (Fig. 2b).

**Fig. 2.**
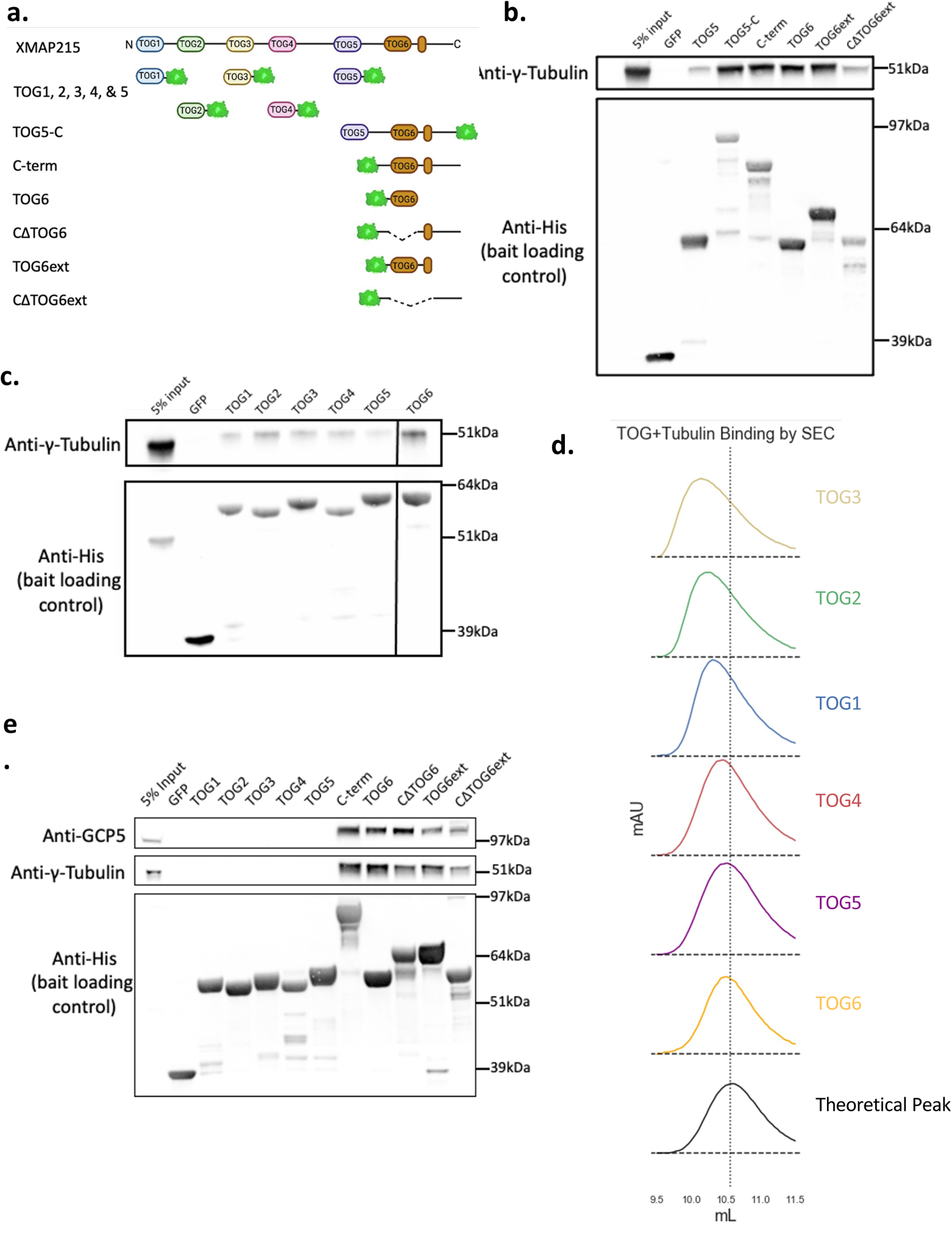
TOG6 specifically binds γ-tubulin and is sufficient to bind γ-TuRC. **a)** Schematic of full length XMAP215 and GFP-tagged constructs used in B-E. Created with Biorender.com **b-c)** Western blot analysis of bead samples from 1μM human γ-tubulin pulldowns using saturated anti-GFP resin. This is representative of three replicates performed. **d)** Chromatograms from gel filtration of 5μM TOG domain constructs mixed with 5μM bovine brain tubulin. The theoretical peak (black) was created by first averaging the traces from all TOG domain control runs and adding this average to the trace of the tubulin-alone control (Fig. S2a). Colored peaks represent observed shifts of each TOG+Tubulin binding reaction from individually calculated theoretical peaks (Fig. S2a). **e)** Western blot analysis of bead samples from a *X. laevis* γ-TuRC pulldown using 10ul of peak sucrose gradient fractions (Fig. S2c) and saturated anti-GFP resin. This is representative of three replicates performed, except for TOGs 1, 2, 3, and 4 for which two replicates were performed.

Being able to bind to γ-tubulin is a novel feature of a TOG domain. To ascertain that this is unique to TOG6, we tested whether TOG domains 1 to 5, which are known to bind αβ-tubulin^4,22,30^, for this property. Using the same pull-down assay with GFP-tagged TOG domains as bait, only TOG6 was able to significantly bind to γ-tubulin (Fig. 2c), making this a unique property of TOG6. To assess whether the reverse is also true, we compared each TOG domain’s ability to bind to ɑβ-tubulin in a gel filtration assay, where binding causes a shift to an earlier elution volume compared to the individual proteins. Tubulin strongly bound to a TOG1-5 construct (Fig. S2a), as well as N-terminal TOG domains 1,2, and 3, whereas TOG4 bound to a lesser extent (Fig. 2d, S2a). Interestingly, TOG domains 5 and 6 failed to notably shift the elution peak, indicating their inability to bind soluble tubulin (Fig. 2d, S2a). Thus, while the N-terminal TOG domains recognize ɑβ-tubulin, TOG6 binds specifically to γ-tubulin, and TOG5 does not bind either.

Finally, we sought to clarify whether the interactions we observed with soluble γ-tubulin also apply to the intact *Xenopus leavis* γ-TuRC. As expected, XMAP215’s C-terminal domain pulled down γ-TuRC, as assayed by probing bead samples for γ-TuRC subunits γ-tubulin and GCP5, and TOG6 alone was also sufficient to pull down γ-TuRC. Interestingly, deletion of TOG6 from the C-terminal domain (CΔTOG6) did not fully abrogate the binding to γ-TuRC. This indicates that there is an additional binding site in the C-terminal domain of XMAP215 other than TOG6. A larger deletion, CΔTOG6ext, did appear to reduce this binding to γ-TuRC, suggesting that the C-terminal four helix bundle and/or the intervening linker also contributes to the interaction (Fig. 2e). To conclude, TOG6 specifically binds to γ-tubulin in isolation and within γ-TuRC. Moreover, we deduce that there must be at least one additional γ-TuRC binding site within XMAP215’s C-terminal domain, likely involving the ext domain and a site other than γ-tubulin.

### Binding to γ-TuRC by XMAP215 is required for its function as a co-nucleator

How important are TOG6 and the ext domain for XMAP215’s ability to act as a co-nucleator with γ-TuRC? To address this, we used total internal reflection (TIRF) microscopy and time lapse imaging to capture the nucleation of single microtubules from γ-TuRCs tethered to passivated coverslips in the presence of various C-terminal deletion constructs of XMAP215 (Fig. 3a-b, Movie S1) (Thawani 2020, Consolati 2020, Rale 2022). In comparison to full-length XMAP215, XMAP215 lacking its C-terminal domain (TOG1-5) nucleated roughly 5-fold less. Similarly, the XMAP215ΔTOG6ext construct, which is missing only amino acids comprising TOG6, the ext domain, and the intervening linker, nucleated at a reduced rate (approximately 6-fold slower than the wild-type protein). The XMAP215ΔTOG6 construct, which is missing only the amino acids comprising TOG6, nucleates roughly 3-fold slower than the full-length protein, i.e. at an intermediate level consistent with our pull-down assay (Fig. 3d, S3b). All constructs produce similar microtubule growth rates, and in turn achieved comparable maximum microtubule lengths. Accordingly, plots of microtubule mass over time follow the same trends as plots of microtubule number (Fig. S3c).

**Fig. 3.**
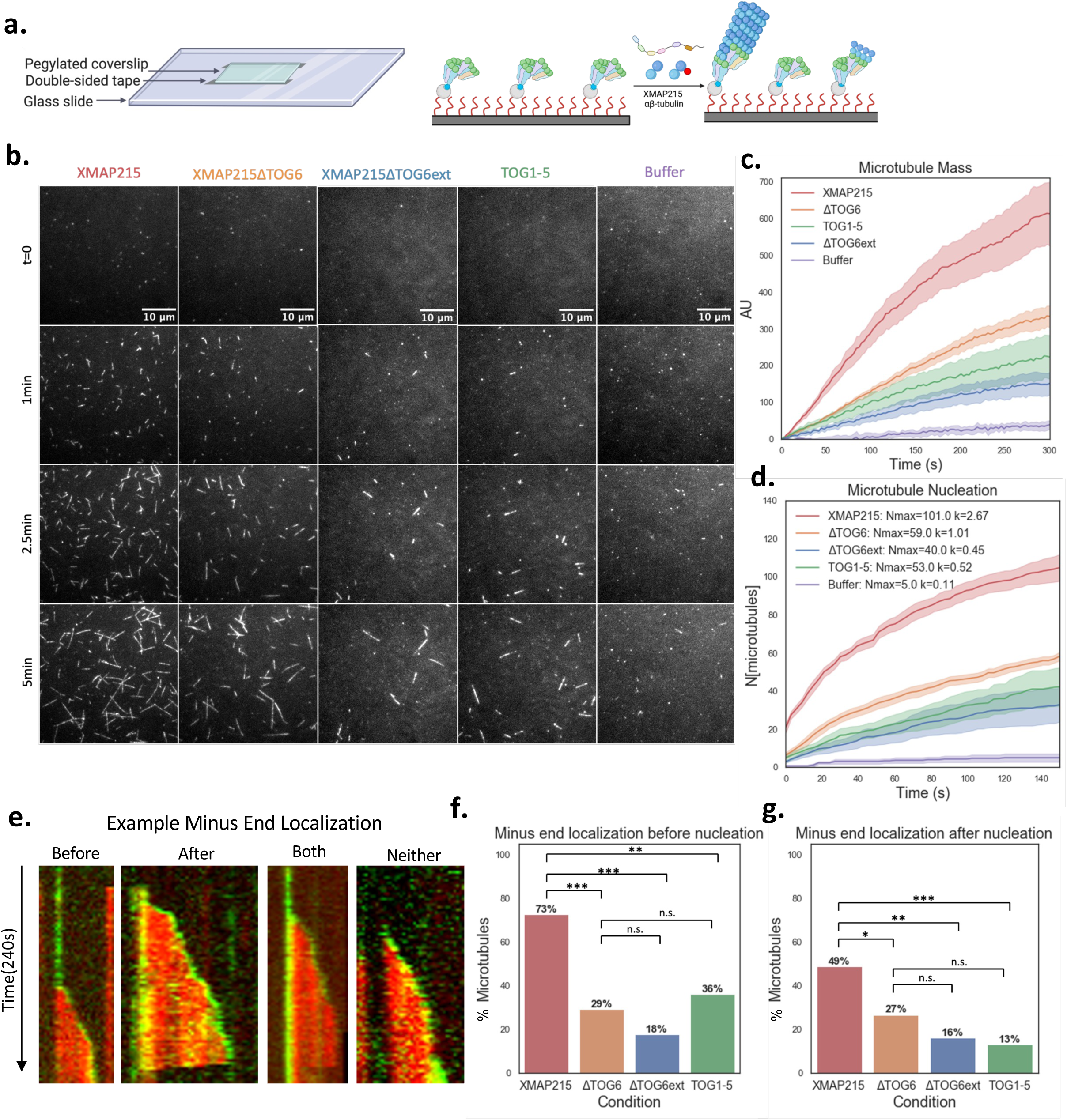
TOG6 and the ext domain promote microtubule nucleation by localizing XMAP215 to γ-TuRC. **a)** Schematic illustrating the setup of our single filament nucleation assays. Passivated biotin-PEG coverslips are used to make flow channels (left), and biotinylated γ-TuRCs are tethered to the coverslips *via* neutravidin (right). Reaction mixes including 7% Alexa Fluor 568 labeled tubulin and XMAP215-GFP constructs or control buffer are flowed in and imaged using time-lapse TIRF microscopy. **b)** Time-lapse images from single filament microtubule nucleation reactions described in A, using 7μM tubulin and 20nM XMAP215-GFP constructs. Microtubules are white against a dark background. **c)** Plot of microtubule intensity over time for the reactions in B. The averages of at least three reactions are plotted with the shaded region representing the corresponding SEM. **d)** Plot of cumulative γ-TuRC nucleated microtubules over time in the reactions from B. The averages of at least three reactions are plotted with the shaded region representing the corresponding SEM. Nmax is the calculated maximum number of microtubules nucleated and k is the calculated nucleation rate (see methods). **e)** Kymographs representing the various microtubule minus-end binding behaviors observed for XMAP215-GFP constructs in the reactions in B. Alexa568-Tubulin is red and XMAP215-GFP is green. **f-g)** Bar plots illustrating the percentage of kymographs exhibiting XMAP215-GFP construct localization at the microtubule minus end before and after microtubule nucleation, respectively, from reactions in B. A proportions Z-test was performed. n.s. represents p≥0.05, * represents 0.05>p>0.01, ** represents 0.01>p>0.001, and *** represents 0.001>p.

We next dissected the role of XMAP215 regions in facilitating microtubule minus end localization. Specifically, localization of XMAP-GFP constructs were visualized as a microtubule was nucleated from γ-TuRC and analyzed via kymographs. We detected four behaviors of XMAP215-GFP constructs with regards to minus end localization (Fig. 3e). (i) GFP localization to the origin of nucleation could occur before tubulin signal was visible then depart as the microtubule began elongating (“before”), (ii) GFP signal could coincide with the appearance of tubulin signal and remain at the minus end following nucleation (“after”), (iii) GFP could localize before tubulin signal appeared and also remain bound after the microtubule began polymerizing (“both”), (iv) and other times no minus end localization was observed (“neither”) (Fig. 3e). We therefore asked whether perturbations of the C-terminal domain would affect our constructs’ localization to the minus end before and after nucleation. Using an anonymized set of kymographs from each condition, we quantified the frequency of each localization behavior (Fig. 3f-g). Indeed, we observed full length XMAP215 localized to the minus end before and after nucleation with frequencies of 73% and 49% respectively, similar to what had been previously reported (Thawani et al., 2018), and that these frequencies were higher than for the C-terminal deletion constructs. Frequency of localization to the minus end after nucleation also appeared to correlate with the size of each C-terminal deletion (Fig. 3g). Taken together, these data suggest that improved γ-TuRC binding by XMAP215 results in faster nucleation rates.

### TOG5 localizes XMAP215 to drive nucleation and polymerization

Albeit to a lesser extent than full length XMAP215, γ-TuRC mediated microtubule nucleation is promoted in the presence of TOG1-5 (Fig. 3a-c, Movie S1). We thus sought to account for how this occurs in the absence of a direct interaction between γ-TuRC and the TOG1-5 construct by examining whether TOG1-5 is sufficient for XMAP215’s activity as an independent nucleation factor. Although nucleation cannot be observed as closely without tethered γ-TuRCs, we used TIRF microscopy to visualize spontaneously nucleated microtubules which fell to the coverslip as a proxy. Over a five minute period, roughly equivalent numbers of microtubules were detected in the presence of either XMAP215 or TOG1-5, and this number was greater than the buffer control (Fig. S4a-b, Movie S3). Thus, XMAP215 can promote spontaneous microtubule nucleation independently of γ-TuRC in vitro as previously shown, and TOG1-5 is sufficient for this activity.

We hypothesized that in order for TOG1-5 to promote nucleation from γ-TuRCs without interacting with it directly, TOG1-5 may interact with the nascent microtubule assembling on γ-TuRC. We generated further XMAP215 truncations, namely TOG1-4 and TOG1-linker4, which contains the linker region previously suggested to promote lattice binding but lacks TOG5, and compared their ability to bind microtubule seeds stabilized with the slowly hydrolysable GTP analog, GMPCPP ^39^. While inclusion of the linker4 microtubule binding domain barely improved the localization of the TOG1-4 construct, TOG1-5 strongly localized to GMPCPP seeds (Fig. S4c-d). This was not due to an avidity effect of TOG domains, because TOG2-5 maintained the same stronger microtubule binding as TOG1-5 while having only four TOG domains, like TOG1-4 and TOG1-linker4. Thus, we uncovered a role of TOG5 in localizing XMAP215 to the microtubule lattice.

How is the ability to bind the MT lattice related to the ability to nucleate a microtubule? We assessed γ-TuRC-dependent microtubule nucleation in the presence of TOG1-5, TOG1-linker4, and TOG1-4. Whereas TOG1-linker4 or TOG1-4 nucleated at similar rates, TOG1-5 stimulated nucleation roughly 3-fold faster (Fig. 4a-c, S5a, Movie S2). Polymerization rates and maximum microtubule length promoted by these constructs reflected the trend observed for nucleation (Fig. S5b). Therefore, the presence of a microtubule binding domain enhances XMAP215’s ability to nucleate microtubules with γ-TuRC, and this microtubule binding activity resides within TOG5.

**Fig. 4.**
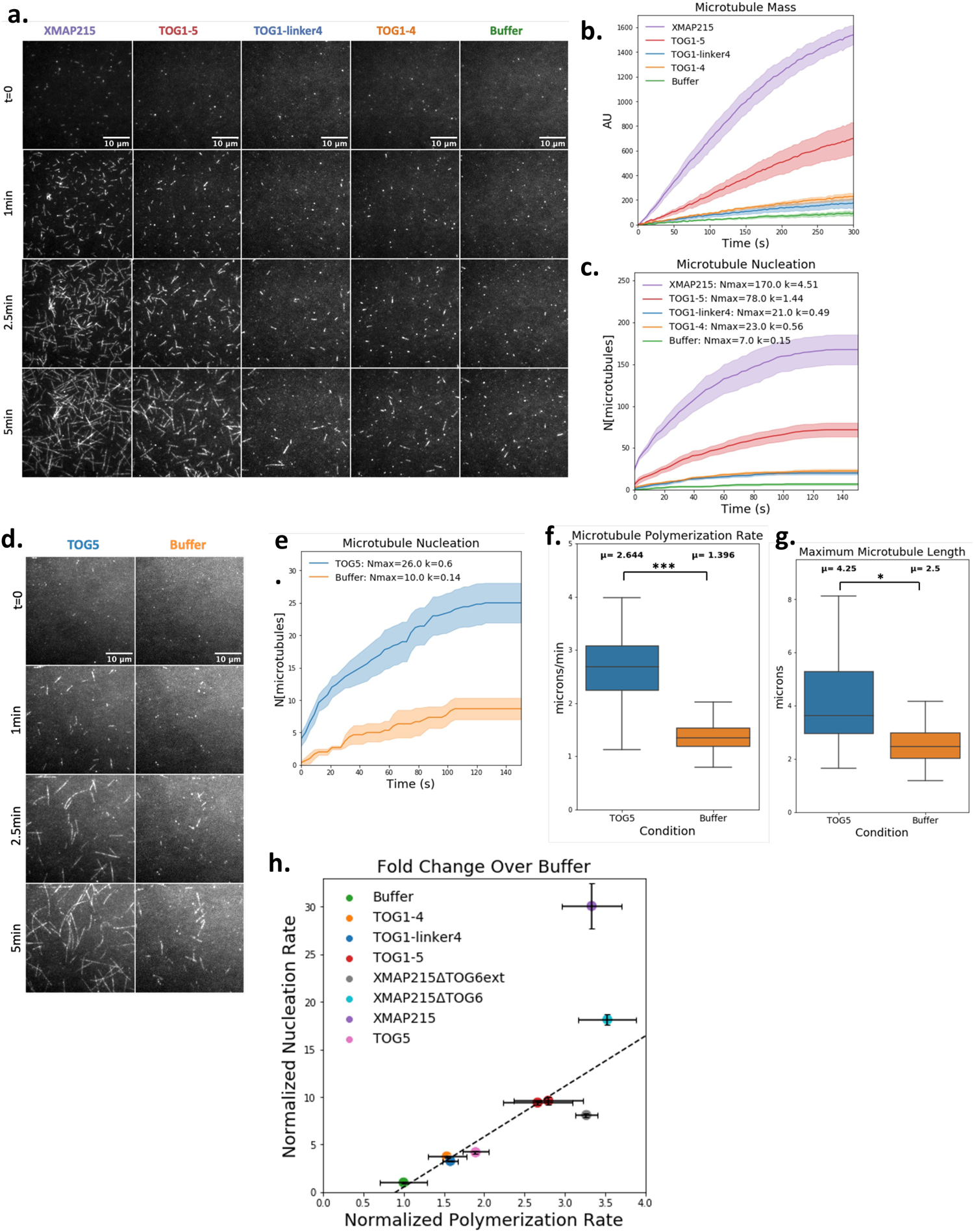
TOG5 promotes microtubule nucleation as part of larger XMAP215 constructs and on its own. **a)** Time-lapse images from single filament microtubule nucleation reactions using 7μM tubulin and 20nM XMAP215-GFP constructs. Microtubules are white against a dark background. **b)** Plot of microtubule intensity over time for the reactions in A. The averages of at least three reactions are plotted with the shaded region representing the corresponding SEM. **c)** Plot of cumulative γ-TuRC nucleated microtubules over time in the reactions from A. The averages of at least three reactions are plotted with the shaded region representing the corresponding SEM. Nmax is the calculated maximum number of microtubules nucleated and k is the calculated nucleation rate (see methods). **d)** Time-lapse images from single filament microtubule nucleation reactions using 10μM tubulin and 1μM TOG5-GFP and a Buffer control. Microtubules are white against a dark background. **e)** Cumulative γ-TuRC nucleated microtubule plot comparable to that in C for TOG5-GFP and buffer reactions from D. **f)** Box and whisker plots of microtubules polymerization rate and **g)** maximum microtubule length for the reactions in D. Plots represent data from at least three reactions. Student’s t-test was performed to compare the mean values calculated from each individual reaction for each condition. μ represents the overall average. n.s. represents p≥0.05, * represents 0.05>p>0.01, ** represents 0.01>p>0.001, and *** represents 0.001>p. **h)** Plot of measured polymerization rates and calculated nucleation rates from single filament γ-TuRC nucleation reactions in the presence of the various XMAP215-GFP constructs normalized to the buffer control. Values from constructs used in Fig. 3 were further normalized according to the full-length XMAP215. Error bars represent one standard deviation derived for each construct from the average polymerization rates in its constituent reactions, and from the fits of the best fit lines in S3B, S5A, and S5E (see methods). The dashed line is a best fit calculated from all constructs except full-length XMAP215, which was excluded as an extreme outlier, y = 4.7478x - 3.9778, R² = 0.9788.

We next sought to identify which TOG domains can individually influence γ-TuRC dependent nucleation. Previous works have established that arrays of multiple TOG domains are necessary for XMAP215 constructs to promote microtubule polymerization^40^. Because individual TOG domains should therefore lack the canonical polymerase activity of wild-type XMAP215, we used a higher concentration of tubulin (10 uM) to bias microtubule dynamic instability more towards growth and added each construct at the high concentration of 1 uM so that we might detect subtle effects on microtubule assembly. TOG6 did not appreciably affect generation of microtubule mass when compared to the buffer condition (Fig. S5c-d, Movie S4). Thus, TOG6 binds γ-TuRC directly but does not change γ-TuRC’s conformation to induce nucleation. TOG1 and TOG4 also had no effect (Fig. 5c-d, Movie S4); however, TOG5 stimulated microtubule nucleation via γ-TuRC about 4-fold (Fig. 4d-e, S5c-e, Movie S4). Remarkably, TOG5 also exhibited plus end tracking behavior and increased the microtubule polymerization rate (Fig. 4e, S6a). This suggests that not only does XMAP215 track the plus end through a ratcheting mechanism involving serial incorporation of tubulin dimers^29,40,41^ but specifically recognizes the tip-specific GTP lattice through TOG5. The increase in polymerization rates further suggests that TOG5 binding promotes lattice formation at growing microtubule ends, and similar activities at the γ-TuRC template should accordingly promote nucleation.

**Fig. 5.**
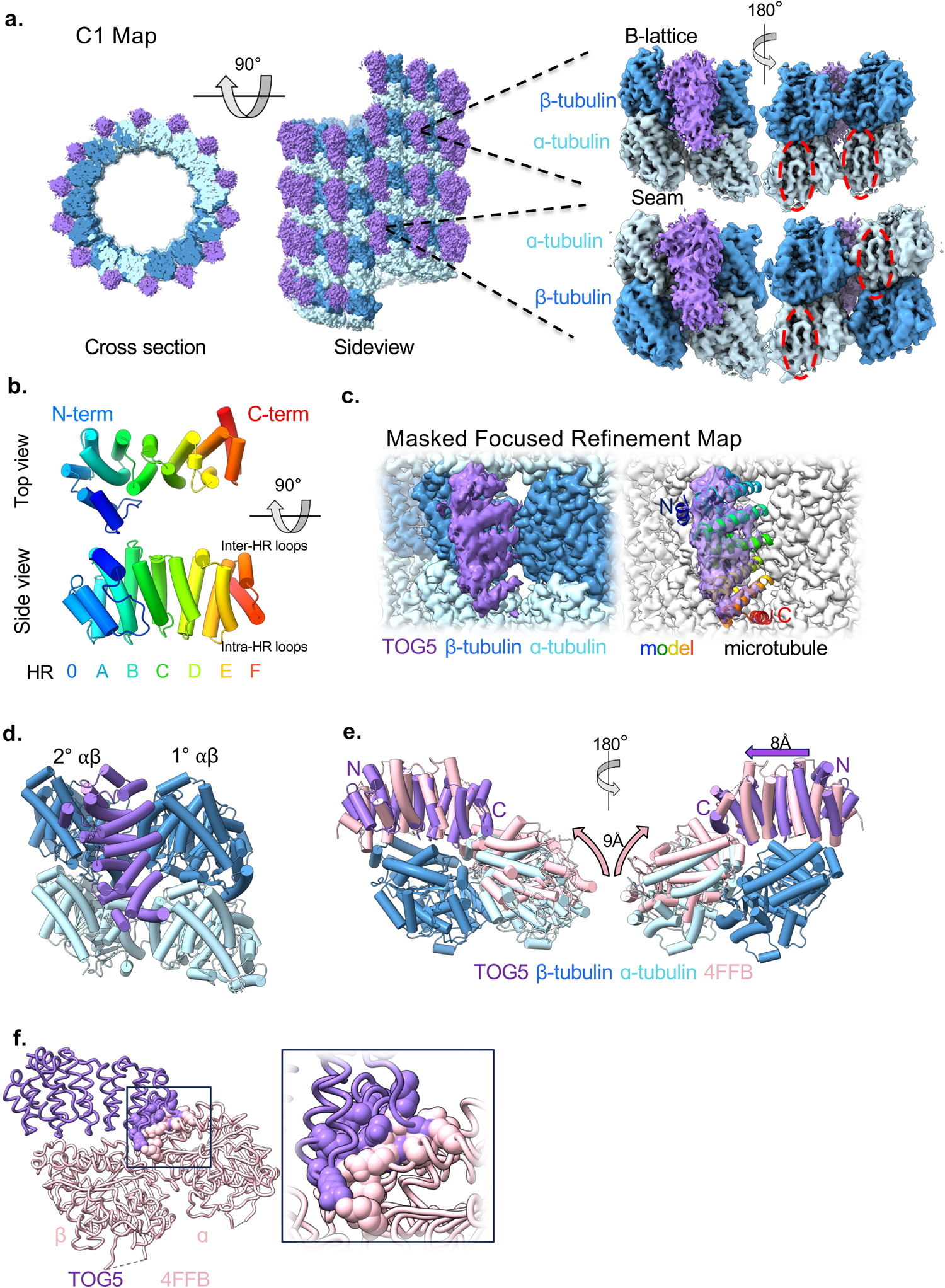
Structure of TOG5 bound to the GMPCPP microtubule. **a)** C1 reconstruction of the TOG5 decorated GMPCPP microtubule. TOG5 density is in purple, ɑ-tubulin in light blue, and β-tubulin in dark blue. Leftmost is a cross section of the reconstruction, the face of which is shown immediately to the right with the microtubule seam centered and the plus end pointed upwards. Dashed lines point towards cropped portions of the C1 map from the Β-lattice (top right) and seam (bottom right). Here, the s9-s10 loops are circled in dashed red. **b)** AlphaFold2 prediction of XMAP215 TOG5 colored from N-terminus (blue) to C-terminus (red). The top view shows the non-tubulin binding surface. Constituent HEAT repeats (HRs) labels are colored correspondingly with the model. **c)** A focused refinement map generated using a mask around two tubulin dimers and the TOG5 density at a location opposite the microtubule seam. Left, density is colored by constituent monomer, right TOG5 density is purple, microtubule density is in light gray, and the rainbow AlphaFold2 TOG5 model is docked into the corresponding density. **d)** Model of TOG5 bound to two GMPCPP lattice incorporated tubulin dimers. The primary and neighboring secondary tubulin dimers bound are indicated 1° ⍺β and 2° ⍺β respectively. **e)** Model from D, secondary tubulin dimer not shown, superimposed with Stu2 TOG1 bound to curved tubulin shown in pink (PDB 4FFB). Models are aligned on the β-tubulin subunit. Pink arrows show displacement of the 4FFB ɑ-tubulin subunit relative to that from our model. The purple arrow shows displacement of TOG5 helices relative to those of TOG1. N- and C-termini of TOG5 are labeled N and C. **f)** Licorice model of TOG5 and 4FFB ⍺β-tubulin from E, with clashing residues shown as spheres. Inset shows magnification of clashing residues.

### Structure of TOG5 bound to the GMPCPP microtubule lattice

How does TOG5 bind the microtubule lattice to promote nucleation and polymerization? To address this, we used single particle cryo-electron microscopy (cryo-EM) to uncover the structure of TOG5 bound to a microtubule. First, we applied microtubule seeds stabilized with GMPCPP to holey carbon grids. We then washed them twice with highly concentrated TOG5-GFP having been desalted into EM buffer (see methods) in order to promote saturation of the microtubule lattice (Fig. S7a). After data collection, we applied a seam-search protocol for accurate particle alignment during processing, then generated a C1 reconstruction of the microtubule (Fig. 5a)^42,43^. The resulting 3.33 Å reconstruction (Fig. S7b) revealed the familiar density of the GMPCPP tubulin lattice in which ɑ- and β-tubulin can be distinguished by the microtubule luminal S9-S10 loop, which is longer in ɑ-tubulin. Strikingly, we observed additional density corresponding with TOG5 which decorated the entire lattice as well as the microtubule seam (Fig. 5a).

We next sought to better resolve the TOG5 density through further processing. We performed symmetry expansion using the measured helical parameters in our C1 reconstruction, then sorted the expanded particles according to TOG5 occupancy using supervised classification, and used a mask for two tubulin dimers plus the TOG5 density for local refinement. Tubulin side chains and helices consistent with TOG5 were visible in the resulting 2.89 Å map (Fig. 5b-c, S7b-c). These were sufficient to build two tubulin dimers and unambiguously dock the AlphaFold2 model of XMAP215 TOG5 for coordinate refinement (Fig. 5c-d). The resulting model constitutes the first structure of a TOG domain bound to the microtubule lattice. The microtubule exterior is recognized by the TOG5’s intra-HEAT repeat loops (Fig. 5b-d), a binding mode consistent with that observed for TOGs 1 and 2 from yeast orthologs (Stu2 and Alp14) bound to soluble tubulin (Ayaz et al., 2012, Ayaz et al., 2014, Nithianantham et al., 2018). This binding occurs primarily to a single tubulin dimer, burying 530 Å^2^ of the β-tubulin surface and 420 Å^2^ of the ɑ-tubulin surface, roughly the same magnitude observed for other TOG/tubulin structures (Fig. S8a). Importantly, TOG5’s unique HR0 buries 100 Å^2^ of a neighboring β-tubulin surface orthogonal to the primary binding site, resembling previous predictions^26^.

To gain atomistic insight into TOG5’s interaction with ɑβ-tubulin and compare its binding mode with other TOG/tubulin interactions, we searched for tubulin side chains within 5Å of TOG5 residues. The search yielded several highly conserved TOG5 residues in common between our structure and an AlphaFold2 multimer prediction of a TOG5/ɑβ-tubulin complex (Fig. S7d-g, S8b). Notably, Phe1250 in the TOG5 HRA loop packs against the backbone in the ɑ11’-ɑ12 loop of β-tubulin (Fig. S7i). This contrasts with the analogous tryptophan residue which occurs in the other canonical XMAP215 TOG domains and binds the β-tubulin ɑ3 helix in TOG1 and TOG2 structures (Fig. S7i)^21,23,24,29^. This altered binding site may help to explain the divergence of this residue in TOG5 from Trp to Phe across orthologs. The second helix of TOG5 HRF also contributes dramatically to the interaction with ɑ-tubulin, accounting for 80% of the interface with TOG5 (330 of 420 Å^2^). This is mediated by TOG5 Arg1452 and Arg1455 flanking Glu414 of ɑ-tubulin, an electrostatic interaction between TOG5 Glu1443 and ɑ-tubulin Arg56, and additional H-bonding and hydrophobic packing by e.g. TOG5 Lys1444 and Met1448, respectively (Fig. S7l-m). Lattice recognition is also surely promoted by TOG5’s unique HR0 loop which forms a lateral contact across tubulin dimers. This interaction is partially electrostatic, involving TOG5 R1222 and a secondary β-tubulin E376 (Fig. S7h). Finally, the HRB loop contributes to primary β-tubulin binding through H-bonding involving TOG5 Thr1289, Thr1291, and Ser1292 (Fig. S7j), and the HRE loop straddles the ɑ-tubulin/β-tubulin interface (Fig. S7k).

In examining our cryoEM reconstructions, we noticed additional density near TOG5 which was positioned consistently with the C-terminal tubulin tails (Fig. S7n). These disordered tails are rich in glutamate residues giving them a negative charge. By plotting Coulombic potential on the surface of our TOG5 model, we see that the tails are revealed at low contour near patches of positive charge on TOG5. This is consistent with XMAP215 having lower affinity for subtilisin treated microtubules^19^. Nonspecific electrostatic interactions with tubulin tails are therefore likely contributing to TOG5 microtubule localization in addition to specific sidechain interactions.

Having gained greater insight into how TOG5 interacts with tubulin we wondered how it is able to discriminate between tubulin’s straight, lattice conformation and its bent, soluble conformation. We aligned TOG1/tubulin structure (PDB 4FFB) from the yeast XMAP215 ortholog Stu2 to our model using the β-tubulin subunits as alignment anchors (Fig. 5e). In doing so, the differences between the bent and straight conformations of tubulin become clear, where the ɑ-tubulin subunit curves up towards the TOG domain by about 9 Å. Further, we noticed that while the first helix of each HEAT repeat aligns fairly well between TOG1 and TOG5 (Fig. 5e left), the second helix of each HEAT repeat in TOG5 is angled or shifted in the direction of ɑ-tubulin for a total displacement of about 8Å (Fig. 5e right). The cumulative effect of these displacements is a severe clash between TOG5 HRE-F and the bent ɑ-tubulin in our aligned model (Fig. 5f). Analogously, TOG1 interacts less with straight ɑ-tubulin (90 Å^2^ buried area) versus bent ɑ-tubulin (350 Å^2^ buried area). Thus, the features of TOG5 that allow it to bind ɑ- and β-tubulin in the straight conformation preclude it from binding bent tubulin, and the features that allow e.g. TOG1 to bind bent tubulin prevent it from strongly binding both monomers in lattice tubulin.

### Binding to γ-TuRC is required for specific nucleation versus polymerization

An outstanding question in the field is what differentiates microtubule nucleation from polymerization^44^. By directly calculating microtubule nucleation and polymerization rates, we report that all XMAP215 truncation constructs that lack the C-terminal domain and thus do not bind γ-TuRC, including TOG5 alone, display a microtubule polymerization rate that is directly proportional to their nucleation rate (Fig. 5h). Strikingly, this linear relationship is broken by the full-length XMAP215 protein. Instead of stimulating nucleation 12.8-fold faster than buffer, according to this linear relationship, we measured an increase of 30.1 ± 2.40-fold. Thus, the ability to directly bind to γ-TuRC is specifically linked to promotion of microtubule nucleation.

We also note that XMAP215ΔTOG6 appears to nucleate slightly better than the trend predicts (18.1 ± 0.6-fold increase versus a predicted 13.8-fold increase in nucleation). This is consistent with our binding data, which show that there is a second binding side other than TOG6, leading to the observed remaining synergy between the XMAP215ΔTOG6 and γ-TuRC. Conversely, XMAP215ΔTOG6ext nucleates less efficiently than predicted by the linear trend (only 8.1 ± 0.2-fold versus the predicted 12.5-fold increase). Interestingly, this is similar to the rate of nucleation of the TOG1-5 construct (9.4 ± 0.4 fold increase), consistent with their similar polymerization rates and their absence of binding to γ-TuRC.

## DISCUSSION

XMAP215 has been well studied as a microtubule polymerase: the unique features and conserved positions of each TOG domain in the pentameric array in vertebrate XMAP215 orthologs were proposed to orient XMAP215 on the growing microtubule to sequentially recruit soluble tubulin, aid its lattice incorporation, and stabilize the newly formed lattice^31^. In this work, we reveal how XMAP215 facilitates microtubule nucleation and discuss its implications below.

Curiously, while genetic duplication events seem to have given rise to homologous TOG domain pairs, TOG1-2 and TOG3-4, TOG5 has remained without a partner^45^. While some NMR characterization of a sixth TOG domain was performed^36,37^, no experimentally validated structural models have been published. Here, we obtained crystallographic evidence for the predicted structure of the TOG6 domain that is conserved across orthologs. This domain is unique in its ability to bind to γ-tubulin and is part of a larger interaction between XMAP215’s C-terminal region and γ-TuRC. The larger interaction involves a four-helix bundle just C-terminal of TOG6, which is also conserved across species. Together, these domains help localize XMAP215 to γ-TuRC for cooperative microtubule nucleation. Thus, we propose that the use of “pentameric” TOG array be revised to “hexameric” TOG array for XMAP215 and its homologues, with TOG5 finally having found its partner domain.

The question remains, how does TOG6 interact with γ-tubulin? To address this, we examined how TOG1/2 and TOG6 could bind to γ-tubulin by performing structural alignments in the following order. First, we aligned beta-tubulin from a tubulin-bound Stu2 TOG2 structure (PDB 4U3J) to γ-tubulin from *Xenopus* γ-TuRC^46^. Then, we aligned *Xenopus* TOG1 and TOG6 to the Stu2 TOG2 according to the least variable heat repeats A-C for TOG1 and heat repeats B-C for TOG6 ^23,24,26^. Were TOG1 to bind γ-tubulin analogously to how it binds b-tubulin, HEAT repeats E-F would clash with the supporting GCP2 (Fig. 6a-b). This may explain why the N-terminal TOG domains exhibited some binding to γ-tubulin but did not bind γ-TuRC. Critically, the kink in TOG6 between HRC and D, where the alpha-solenoid twist transitions from right-handed to left-handed, helps to avoid this clash. Thus, it may be that more efficient nucleation from γ-TuRC through a direct interaction with XMAP215 drove evolution of TOG6’s divergent solenoid architecture.

**Fig. 6.**
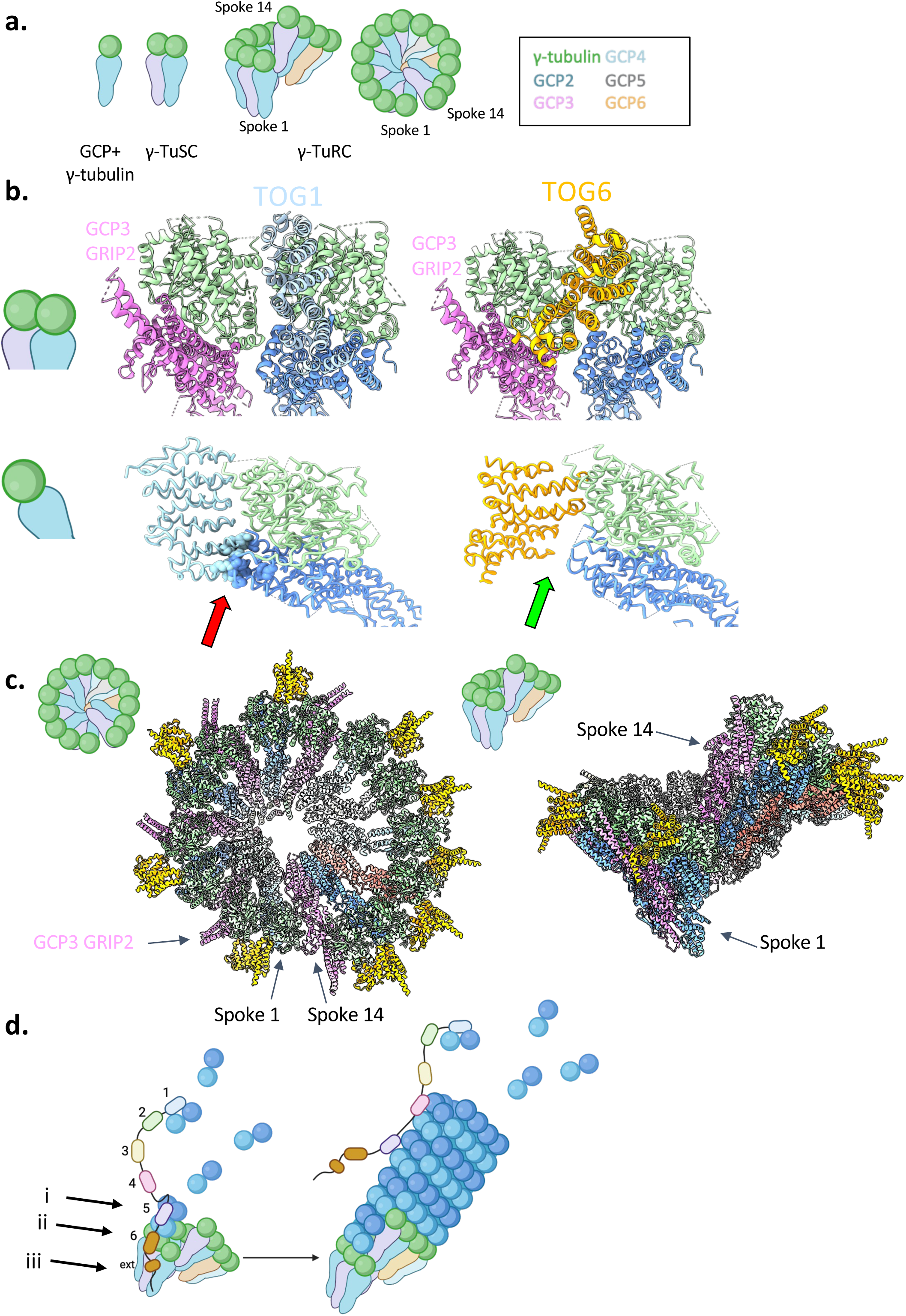
XMAP215 may recognize specific spokes of γ-TuRC to promote microtubule nucleation. All panels of this figure were at least in part created with Biorender.com. **a)** Schematic of γ-TuRC and some of its constituent sub-complexes. **b)** Models of how XMAP215 TOG1 and TOG6 may interact with γ-tubulin as part of the γ-TuSC sub-complex. Top depicts a ribbon model of the TOGs’ modeled interaction with the γ-TuSC from a perspective outside the γ-TuRC ring. Bottom depicts a side view of licorice model with each TOG and GCP2. Clashing residues are shown as spheres and indicated by a red arrow, and the absence of such a clash is indicated by a green arrow. **c)** Models of how XMAP215 could interact with γ-TuRC spokes 9-12 by TOG6 interaction with γ-tubulin. **d)** Schematic representing a model for the activity of the XMAP215/γ-TuRC nucleation module. i) TOG5 binds tubulin assembling on γ-TuRC, ii) TOG6 binds γ-tubulin, and iii) an additional interaction between the XMAP215 C-terminal region and γ-TuRC occurs. TOG domains are labeled on the left with numbers, and the C-terminal four helix bundle is likewise labeled “ext.”

It will further be interesting to explore which spokes of γ-TuRC XMAP215 binds to. The alpha-helical extensions of the GCP3 GRIP2 domain obstruct the probable binding site between TOG6 and γ-tubulin, but this site remains free at GCP2, 4, 5, and 6 (Fig. 6b-c). These 9 available sites are distributed on γ-TuRC in an interesting way, namely at every other spoke in the case of the more symmetric g-TuSC portion (spokes 1-8), and/or at every spoke on the metazoan-specific portion (spokes 9-14, Fig. 6c)^9–11^. It may be that XMAP215 can bind all 9 of these available sites, or that specific interactions with GCP and the small γ-TuRC MTZ subunits ^47^ bias it towards one region of the ring over another.

We further uncovered a role for TOG5 in γ-TuRC dependent microtubule nucleation and described the structural basis for how XMAP215 binds the microtubule. TOG5 is the primary driver of XMAP215’s localization to microtubules, can independently track the microtubule plus end, and accelerate microtubule polymerization and nucleation from γ-TuRC. Our cryoEM structure reveals that TOG5’s unique architecture enables it to bind the microtubule, sensing tubulin curvature within the lattice with its C-terminal HRs E-F, and confirms that its characteristic N-terminal HR0 also contributes a minor interaction with the b-tubulin of a laterally associated dimer. Taken together, we conclude that TOG5 helps to direct XMAP215 to the nascent or growing microtubule plus-end through specific recognition of straight GTP tubulin and propose that TOG5 binding promotes assembly of the microtubule lattice, possibly by stabilizing the straight tubulin conformation, decreasing tubulin off-rates or helping to “zipper up” neighboring protofilaments. It may be that the coupled activities of TOG5 and TOG6 help tether microtubules to, for example, centrosomal γ-TuRCs^5^, and TOG5’s unique lattice binding may also underlie XMAP215’s recently characterized cross linking of the actin and microtubule cytoskeletons^48^. It will further be interesting to structurally compare how CLASPs, another family of TOG domain proteins that stabilize microtubules, interact with microtubules^21^.

There has been an ongoing debate about how microtubule nucleation and polymerization relate to one another. Directly measuring individual microtubule nucleation events in real time and comparing them with polymerization rates allowed us to better understand this relationship. By plotting the fold change in polymerization rate versus fold change in nucleation rate, we revealed a linear trend for XMAP215 constructs which do not bind γ-TuRC. An analogous relationship was previously reported *in vitro* using microtubule seeds and in cells in which tubulin sequestration slowed both nucleation and polymerization^49^. Thus, proteins which aid in the association of GTP tubulin with the microtubule plus end proportionally affect rates of both nucleation and polymerization. However, nucleation activity specifically is promoted as soon as XMAP215 and γ-TuRC directly interact with one another. In fact, increased nucleation efficiency naturally depends on the extent of XMAP215’s interaction with γ-TuRC. Thus, this interaction defines the molecular basis for the cooperativity between XMAP215 and γ-TuRC as the universal nucleation module of the cell.

Taken together, we propose the following model for microtubule nucleation. XMAP215 and γ-TuRC interact directly via two modes, the g-tubulin specific TOG6 domain and the C-terminal region with its four helix bundle which may interact with any of the GCP or MTZ subunits. This direct binding gives rise to the synergy between the two nucleators. The N-terminal TOG5 domain is positioned to interact with the first tubulin dimers assembling onto the γ-tubulin ring template and stabilizes their lattice interactions and straight conformation. Therefore, both γ-TuRC and TOG5 together enable lateral tubulin interactions, which promotes γ-TuRC closure and a switch from nucleation to polymerization. The N-terminal TOG domains 1-4 recruit soluble tubulin to the nascent microtubule lattice. Thus, the unique features and conserved positions of each TOG domain in the hexameric array in vertebrate XMAP215 orthologs orient XMAP215 to recruit soluble tubulin, aid its lattice incorporation, and stabilize the lattice to promote both microtubule nucleation and polymerization (Fig. 6d).

## METHODS

### Protein cloning, expression, and purification

#### XMAP215 truncation constructs

TOG1-5 (1-1,460), TOG1-4 (1-1,090), TOG5-C (1,191-2,065) were generated for a previously published work by subcloning into an ampicillin resistant pST50 vector with C-terminal GFP, 7xHis-StrepII tags (Tan et al., 2005, Thawani et al., 2018). For this work, TOG1(1-240), TOG2 (264-515), TOG3 (556-808), TOG4 (846-1,090), TOG5 (1,191-1,460), TOG1-linker4 (1-1,192), and TOG2-5 (264-1,460) were subcloned into the same pST50 with C-terminal GFP, 7xHis-StrepII tags. XMAP215 C-term (1,461-2,065), CΔTOG6 (1,461-1,559+1,840-2,065), CΔTOG6ext (1,461-1,559+1,932-2,065) were subcloned into a similar pST50 vector, but with N-terminal StrepII-6xHis and GFP tags. TOG6 (1,560-1,839) and TOG6ext (1,560-1,932) were subcloned into a pET-28a vector with N-terminal 6xHis, GFP, HRV3C, and StrepIII tags. For crystallization, TOG6 (1,560-1,839) was subcloned into the ampicillin resistant pST50 vector with an N-terminal 7xHis tag.

All constructs were transformed into Rosetta 2 (DE3) pLysS *E. coli* cells for expression. Cells were grown under antibiotic selection with chloramphenicol and either ampicillin or kanamycin in 1-2L Luria Broth cultures at 37℃ to an optical density (OD) at 595 nm of 0.6-0.8. Cultures were then chilled on ice at 4℃ before induction with 1mM isopropyl β-D-1-thiogalactopyranoside (IPTG), whereupon they were incubated for 16-18 hours at 16℃. Cultures were then pelleted, flash frozen, and stored at −80℃ until they were thawed for purification. Unless otherwise specified, all protein handling during and after purification was performed on ice or at 4℃.

For purification, cell pellets were thawed, resuspended in Strep Binding Buffer (280mM NaCl, 2mM MgCl2, 100mM Tris, 6mM beta-mercaptoethanol (BME), .2mM phenylmethanesulfonyl fluoride (PMSF), pH 7.7) supplemented with 1µg of DNase from bovine pancreas and 1 dissolved tablet of cOmplete EDTA-free Protease Inhibitor Cocktail (Roche). Pellets were then homogenized with a Biospec Tissue Tearor (Dremel), and lysed with an Emulsiflex C3 (Avestin) at 10,000-15,000 psi. Lysates were clarified by ultracentrifugation for 35 minutes at 4℃ at 70,000 x (g) in a 45Ti rotor (Beckman Coulter). Constructs containing N-terminal TOG domains (TOG1, TOG2, TOG2, TOG4, TOG5, TOG1-5, TOG1-linker4, TOG1-4, and TOG5-C) were subjected to affinity purification using StrepTactin SuperFlow resin (IBA) using Strep Binding Buffer (280mM NaCl, 2mM MgCl2, 100mM Tris, 6mM BME, .2mM PMSF, pH 7.7) for lysis, binding, and washing. Elution was performed in Strep Binding Buffer supplemented with 3.3mM desthiobiotin. TOG1, TOG2, TOG3, TOG4, and TOG5 were then dialyzed overnight in gel filtration buffer (10mM HEPES, 100mM KCl, 1mM MgCl_2_, 0.5mM ethylene glycol-bis(β-aminoethyl ether)-N,N,N’,N’-tetraacetic acid (EGTA), 10% sucrose, 6mM BME, 0.2mM PMSF, pH 7.5). TOG1-5 and TOG1-linker4 were further subjected to gel filtration on a Superose 6 Increase 10/300 GL column (Cytiva), and TOG2-5, TOG1-4, and TOG5-C on a Superdex 200 Increase 10/300 GL column (Cytiva) in gel filtration buffer. Peak elution fractions from affinity and size exclusion chromatography were analyzed for purity by SDS-PAGE and staining with Coomassie blue. The desired fractions’ concentrations were estimated using a Bradford assay against a bovine serum albumin (BSA) standard before aliquoting and flash freezing in liquid nitrogen before storage at −80℃.

C-term, CΔTOG6, CΔTOG6ext, TOG6, TOG6ext, and the TOG6 crystallization construct pellets were similarly resuspended, homogenized, and lysed, but instead using His Binding Buffer (300mM NaCl, 50mM NaPO4H, 10mM Imidazole, 6mM BME, .2mM PMSF, pH7.9) supplemented with 1µg of DNAse from bovine pancreas and 1 dissolved tablet of cOmplete EDTA-free Protease Inhibitor Cocktail (Roche). Lysates were incubated with Ni-NTA agarose (Qiagen), washed with His Wash Buffer (His Binding Buffer with 40mM imidazole), and eluted with His Elution Buffer (His Binding Buffer with 250mM imidazole). All constructs were subject to gel filtration on a Superdex 200 Increase 10/300 GL column in gel filtration buffer, except the crystallization construct which was run on a Superdex 75 10/300 GL column (Cytiva) in HBS (10mM HEPES, 100mM NaCl, 1mM dithiothreitol (DTT), pH 7.5). These were similarly evaluated by SDS-PAGE, Coomassie staining, and Bradford assay concentration quantitation before aliquoting, snap freezing, and storage at −80℃.

#### XMAP215 full length, TOG6, and TOG6ext deletion constructs

XMAP215ΔTOG6 (1-1,559+1,840-2,065) and XMAP215ΔTOG6ext (1-1,559+1,932-2,065) were created as derivatives of a pFastBac1 vector containing full-length XMAP215 (1-2,065) with C-terminal TEV, GFP, 7xHis, and StrepII (Thawani et al., 2020) using the Q5 Site Directed Mutagenesis Kit (NEB). All three constructs were expressed using the Bac-to-Bac Sf9 cell expression system (Invitrogen). Full-length XMAP215 was purified from a 3L pellet, and XMAP215ΔTOG6 and XMAP215ΔTOG6ext were each purified from 1L pellets, all of which were snap frozen and stored at −80℃. A similar purification scheme to that described above was used. Briefly, cell pellets were resuspended in Strep Binding Buffer plus DNAse and protease inhibitor cocktail, homogenized by pipetting, and lysed using the Emulsiflex. Lysates were clarified by ultracentrifugation at 4℃ at 184,000 x (g) in a 70Ti rotor for 50 minutes (Beckman Coulter). After affinity chromatography with StrepTactin SuperFlow resin (IBA), proteins underwent gel filtration in gel filtration buffer on a Superose 6 Increase 10/300 GL column, and peak fractions were evaluated by SDS-PAGE, Coomassie staining, and Bradford assay to determine concentration. Peak fractions were aliquoted and snap frozen before storage at −80℃.

#### γ-Tubulin and γ-TuRC

Human γ-tubulin with C-terminal TEV, StrepII, 7xHis tags was expressed and purified as described previously, using the Bac-to-Bac Sf9 expression system, nickel resin, and gel filtration^4^. Endogenous *X. laevis* γ-TuRC was also purified according to a published method^13^. Briefly, a human γ-TuNA (γ-TuRC nucleation activating) peptide from CDK5RAP2 with N-terminal StrepII, 6xHis, TEV, HaloTag, and HRV3C tags was expressed from a pST50 vector in Rosetta2 (DE3) pLysS *E. coli* in 2L of Terrific Broth and purified using StrepTactin Superflow resin. Halo-γ-TuNA was then coupled to Halo Magne beads (Promega) and used to pull endogenous γ-TuRC out from *X. laevis* oocyte extract. Preps for use in microtubule nucleation assays were biotinylated by incubation with 40 µM EZ-Link NHS-PEG4-Biotin for 1 hour (ThermoFisher). γ-TuNA/γ-TuRC was eluted by protease cleavage of the HRV3C site with PreScission overnight. The eluent was then applied to a 10-50% sucrose gradient in CSF-XB (10mM HEPES, 100mM KCl, 2mM MgCl_2_, 0.1mM CaCl_2_, 5mM EGTA, 6mM BME, pH 7.7) plus 0.1mM GTP and ultracentrifuged for 3 hours at 200,000 x g in a TLS55 rotor (Backman Coulter). Sucrose gradients were manually fractionated and evaluated using a 4-12% Bis-Tris SDS-PAGE gel and subsequent SNAP I.D. using an anti γ-tubulin GTU-88 antibody (Sigma). The peak fraction (either 7 or 8) was then aliquoted and flash frozen.

### X-ray crystallography

#### Crystallization

A construct of *X. laevis* XMAP215 TOG6 (aa1563-1839) was crystallized by hanging-drop vapor diffusion at 20°C after mixing 1 μL of protein (7.5 mg/mL) with 1 μL of well buffer (75 mM Tris, 25 mM MES, 600 mM NaCl, 4% PEG 8,000 (w/v), 30% Ethylene Glycol (v/v), pH 6.75). Crystals exhibited needle morphology with a hexagonal cross section and were roughly 0.3-0.4 x 0.1 x 0.1 mm and grew within a week. Crystals were flash frozen in liquid nitrogen without need for additional cryoprotectant. X-ray diffraction data were collected at the U.S. National Synchrotron Light Source II AMX Beamline.

#### Molecular Replacement and Refinement

Data were processed using XDS. The AlphaFold2 prediction for the crystallized construct was then used for molecular replacement in PHASER. Successful molecular replacement required *in silico* deletion of the first alpha-helix of the first heat repeat and therefore did not appear in the resulting map, but subsequent attempts to purify a construct without this helix revealed greatly decreased its solubility and resulted in the majority of protein precipitating during purification. Further rigid body refinement was performed using LORESTR.

### Protein binding assays

#### GFP Pulldowns

Pulldowns were performed using GFP-Trap Magnetic Agarose Resin (Chromotek) for each reaction. Resin was first equilibrated in bulk with three 1mL washes of gel filtration buffer (10mM HEPES, 1mM MgCl2, 1mM EGTA, 100mM KCl, 6mM BME, 10% sucrose (w/v), pH 7.5), then aliquoted such that 5μL of packed resin could be used for each pull down. 100μL of each GFP-tagged bait construct in gel filtration buffer was then coupled to the equilibrated resin rotating for 1 hour. To ensure saturation, sufficient amounts of each bait construct were used to have 40μg equivalents of GFP present, twice the capacity for 5μL of resin. The resin was then washed once with 200μL of gel filtration buffer, then resuspended in 200μL blocking buffer (10mM HEPES, 5mM MgCl2, 1mM EGTA, 100mM KCl, 6mM BME, 2% sucrose (w/v), 0.05% Tween (v/v), 1% bovine serum albumin (w/v), pH 7.5) and incubated rotating for 3 hours. The blocking buffer was subsequently removed and replaced with 100μL of 1μM Human γ-tubulin in blocking buffer or 90μL blocking buffer plus 10μL peak *Xenopus* γ-TuRC fraction and allowed to bind rotating for 1 hour. Blocking buffer was supplemented with 2mM GTP for these binding reactions. The resin was then washed 3 times with 200μL of wash buffer (10mM HEPES, 5mM MgCl2, 1mM EGTA, 100mM KCl, 6mM BME, 2% sucrose (w/v), 0.05% Tween (v/v), pH 7.5) and finally resuspended in 100μL of 1xSDS-PAGE sample buffer (6mM DTT) and boiled for 10 minutes.

Pull down samples were then run on a 4-12% Bis-Tris denaturing gel (GenScript), then analyzed by Western blotting. First, membranes were probed for GCP5 (sc-365837, Santa Cruz Biotechnology) and/or γ-tubulin (T6557, Sigma), then stripped by short incubation in Western Blot Stripping Buffer (Thermo Scientific), and re-probed for 6xHis (MA1-21315, Invitrogen) to evaluate bait protein coupling.

#### Size exclusion chromatography

For size exclusion assays we used an AKTA PURE SYSTEM with Superdex 75 10/300 GL (Cytiva) column which was equilibrated with BRB80 (80 mM PIPES, 1 mM MgCl2, 1 mM EGTA, pH 6.8) supplemented with 0.05% Tween-20 (v/v), and the same was used to dialyze TOG-GFP constructs. TOG-GFP constructs were dialyzed for 1 hour using a 3,500 MW Slide-A-Lyzer Mini Dialysis Cassette (Thermo Scientific). Bovine brain tubulin (PurSolutions) was diluted 1 to 10 in cold BRB80, then both the tubulin and dialyzed TOG-GFP were separately precleared by ultracentrifugation at 247,000 x g in a TLA100 rotor (Beckman Coulter) for 10 minutes at 2°C. The concentrations of the clarified proteins was estimated by Bradford (BioRad) assay against a bovine serum albumin (BSA) (Sigma) standard curve. 100 μL samples were mixed containing 5 μM TOG-GFP construct, 5 μM tubulin, or both in BRB80 plus 0.05% Tween-20 and 1 mM GTP and incubated on ice for 10 minutes before loading onto the column. Absorbance at 280 nm was used to track the protein peak.

### *In vitro* TIRF assays

#### γ-TuRC mediated microtubule nucleation

These assays were performed as established in previous publications ^10,12^, with slight modifications as described in detail by Rale and coworkers (2023). Briefly, imaging chambers were made with two pieces of parallel double-sided tape on a slide passivated with PLL-g-PEG (SuSOS AG), and by attaching silanized, PEGylated, High Precision 22×22 mm No 1.5H coverslips (Deckglaser) with 10% biotin-PEG (w/w) which had been previously etched with NaOH and Piranha solution. Coverslips were used within three months of preparation and imaging chambers were made fresh.

Before imaging, chambers were blocked with 5% v/v Pluronic F-127 casein buffer (BRB80 with 30 mM KCl, 1 mM GTP, 0.075% w/v methylcellulose 4000 cp, 1% w/v D-glucose, 0.02% w/v Brij-35, 5 mM BME, 0.05mg/mL κ-casein). NeutrAvidin (ThermoFisher) at 0.05 mg/mL in casein buffer was then flowed in and used to attach biotinylated γ-TuRC diluted 1:100 in BRB80 to the coverslips.

Reaction mixes were prepared with the desired ⍺β-tubulin concentrations (7 μM or 10 μM) containing 5% Alexa 568-tubulin plus 1 mg/ml BSA (Sigma) in assay buffer (BRB80 with 30 mM KCl, 1mM GTP, 0.075% w/v methylcellulose 4000 cp, 1% w/v D-glucose, 0.02% w/v Brij-35, 5 mM BME). XMAP215-GFP constructs or control gel filtration buffer (10 mM HEPES, 100 mM KCl, 1 mM MgCl_2_, 0.5 mM EGTA, 10% sucrose, 6 mM BME, pH 7.5) were added to the desired concentration before clarification by ultracentrifugation at 247,000 x (g) for 10 minutes at 2°C in a TLA100 (Beckman Coulter). Glucose oxidase (GmbH) was then added to the clarified mix at 0.68 mg/mL and catalase (Sigma) at 0.16 mg/mL before flowing into the imaging chamber. Imaging was performed on a Nikon Ti-E inverted stand (RRID:SCR_021242) with an Apo TIRF 100 x oil objective (NA = 1.49) and an Andor iXon DU-897 EM-CCD camera with EM gain set to 300, with the objective heated to 33.5°C using a heated collar (model 150819–13, Bioptechs). Imaging began after precisely 1 minute. A single field of view was imaged every 3 seconds for 5 minutes. Microtubule mass was quantitated after first using FIJI/ImageJ’s^50^ “Subtract Background’ function with a rolling ball radius of 50 pixels on all frames, measuring the mean intensity of the tubulin fluorophore for each frame, then subtracting the value of the first frame from all subsequent frames. To measure microtubule nucleation and dynamics, we cropped each movie to the middle 256×256 pixels and 150 seconds. We used the Stackreg or HyperStackReg plugins to correct translational drift^51,52^, then manually selected each microtubule using the tubulin channel only for the “Reslice” operation. The generated kymographs were analyzed to determine the frame in which nucleation occurred, microtubule growth rate, and maximum microtubule length. Microtubules which were not nucleated by tethered γ-TuRCs (i.e. exhibiting two dynamic ends or drift of the entire microtubule) were excluded from our quantification. The microtubule nucleation rate was calculated by fitting equation (1) to the mean cumulative curve of the number of microtubules nucleated over the first 150 seconds for the time-lapses belonging to each condition:

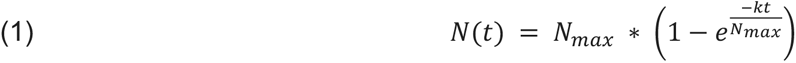

where *N*(*t*) is the number of microtubules nucleated by time t, and *N_max_* is the maximum number of microtubules nucleated. Box plots of microtubule polymerization rates and maximum length were generated using data from each individual microtubule, while nonparametric t-tests were performed using averages for all the microtubules in each movie.

To quantify minus (non-polymerizing) end localization in live γ-TuRC nucleation assays, we generated 30-41 total dual-colored kymographs of random microtubules from separate reactions using each XMAP215-GFP C-terminal deletion construct. All kymographs from across conditions were pooled and randomized before a blind analysis in which we categorized them based on GFP signal on the minus end before and after microtubule nucleation. GFP signal had to be visible at least 3 pixels before Alexa 568 signal appeared to be counted as localizing before nucleation and remain clearly present during the lifetime of the microtubule to be counted as localizing after nucleation.

#### Spontaneous microtubule nucleation

Imaging chambers for spontaneous microtubule nucleation assays were prepared identically to those described in the previous section, but without NeutrAvidin or γ-TuRC steps. Reaction mixes were prepared using 5% Alexa 568 labeled 15 μM tubulin plus XMAP215-GFP construct or control gel filtration buffer, clarified, and imaged using the same microscopy setup as above. We imaged 9 fields of view in a 3 x 3 array every 30 seconds for five minutes using the NIS-Elements AR program (Nikon), and manually counted the number of microtubules in all fields of view in every other frame for plotting.

#### Microtubule localization

To assay binding of constructs to the microtubule lattice, we made Guanosine-5’-[(α,β)-methyleno]triphosphate (GMPCPP, Jenna Bioscience) stabilized microtubule seeds using a standard protocol. A polymerization mix of 20 μM bovine brain tubulin (PurSolutions) containing 10% biotin- and 10% Alexa 568-labeled tubulin was prepared in BRB80 (80 mM PIPES, 1 mM MgCl2, 1mM EGTA, pH 6.8) with 1 mM GMPCPP. This mixture was incubated on ice for 5 minutes before ultracentrifugation at 247,000 x (g) in a TLA100 rotor for 10 minutes at 2°C and subsequent incubation at 37°C for 1 hour. The resulting microtubule seeds were pelleted by centrifugation at 17,000 x g for 15 minutes at room temperature, then resuspended in one reaction volume of warm BRB80 using a wide orifice pipette tip.

Imaging chambers were prepared as for the γ-TuRC mediated nucleation assays, though the PLL-g-PEG step was skipped and GMPCPP seeds diluted 1:500 in warm BRB80 were attached to coverslips in place of γ-TuRC. Reaction mixes were prepared with XMAP215-GFP constructs at 200 nM in localization assay buffer (BRB80 with 75 mM KCl, 1% w/v D-glucose, 5 mM BME, 0.1 mg/mL BSA, 0.34 mg/mL glucose oxidase, 0.08 mg/mL catalase), flowed into imaging chambers, and imaged after a 10 minute incubation at room temperature. Imaging was performed on a Nikon Ti-E inverted stand (RRID:SCR_021242) with an Apo TIRF 100 x oil objective (NA = 1.49) and a Hamamatsu ORCA-Fusion BT Digital CMOS camera. Images were captured in a 3 x 3 array using the NIS-Elements AR program (Nikon).

For processing, we used a custom FIJI/ImageJ macro^50^. To segment microtubules, we automatically thresholded the tubulin channel using the Otsu method, performed the “open” operation followed by the “dilate” operation to remove punctae and encompass all the microtubule signal. We accepted regions of interest (ROI) with a 0.75 μm^2^ minimum particle area. The mean fluorescence intensity in both the tubulin and XMAP215 channels for each ROI was recorded. To account for background fluorescence, we created an inverse mask by again automatically thresholding, dilating once, and inverting the selection to exclude all microtubules. We measured the mean intensity in both channels over the entire remaining area and subtracted the resulting value for each channel from the corresponding values for each segmented microtubule. The difference was used to calculate the MAP/MT values reported. Steps were taken to manually exclude protein aggregates from intensity measurements.

### Cryo-Electron Microscopy

#### Sample Preparation

GMPCPP stabilized microtubule seeds using a slightly modified protocol in which we skipped the microtubule pelleting step. A polymerization mixture containing 20 μM bovine brain tubulin (PurSolutions) in BRB80 (80 mM PIPES, 1 mM MgCl2, 1 mM EGTA, pH 6.8) plus 1 mM GMPCPP was incubated on ice for 5 minutes before preclearing by ultracentrifugation at 247,000 x g for 10 minutes at 2°C. The supernatant was then incubated at 37°C for 1 hour, then finally diluted 1:2 in warm BRB80. Meanwhile, purified TOG5-GFP construct was desalted into EM Buffer (BRB80 plus 5 mM BME and 0.05% Tween-20 (v/v)) using a 0.5 mL 7K MWCO Zeba Spin Desalting Column (Thermo Scientific). After desalting, we estimated the TOG5 concentration to be 66 μM by Bradford Assay. Quantifoil R1.2/1.3 400-square mesh Cu grids (Quantifoil) were glow-discharged for 1 minute at 10 mA using a PELCO easiGLow glow discharging system (Ted Pella) and loaded onto a Leica EM GP2 plunge freezer (Leica Microsystems) with the chamber set to 20°C and 85% humidity. 3 μL of diluted microtubule seeds were applied to the grid, manually blotted away with Whatman No. 1 blot paper, then washed twice with 3 μL of TOG5 in EM buffer. A 30 second incubation took place after each application of microtubules or TOG5 to the grid. The grid was then front blotted for 3 seconds using the Leica EM GP2 and plunge-frozen in liquid ethane and transferred to a grid box under liquid nitrogen where it was stored until data collection.

#### Cryo-EM data collection

Data was collected using a Titan Krios G3 cryo Transmission Electron Microscope (Thermo Fisher Scientific) at 300 kV with a K2 Summit direct electron detector camera equipped with a GIF Bio-Quantum energy filter (Gatan). Collection of 32 frame movies was automated using EPU in EFTEM mode with a 4.1 second exposure time, total dose of 15 e^-^/Å^2^, a pixel size of 1.114 Å (later binned to 1.405Å during processing), and a defocus range of −2 to −3.5 μm. We collected 2,486 movies total.

#### Cryo-EM data processing

The microtubule data processing was done using a similar protocol as previously described ^43^. Briefly, MT segments were automatically picked using ‘Filament Tracer’ in CryoSPARC^53^. The microtubule particles with 8-nm step size were subject to 2D classification and 3D classification (13-protofilaments vs 14-protofilaments) in CryoSPARC. The 14-protofilament microtubule particles were used for subsequent data processing. We used a previously established protocol to determine the seam location for each MT particle^42^, and then did symmetry expansion using the 3-start symmetry (rise 8.9 Å, twist −25.8°) in CryoSPARC, which effectively rotates/translates all the pseudo-helical symmetry related tubulin subunits to the same location for further local 3D classification and local refinement. We performed supervised 3D classification (2 classes) in CryoSPARC to sort out the bound and unbound states of TOG5. Finally, we could achieve 2.9 Å local resolution via local refinement in CryoSPARC using a soft-edged mask covering two tubulin dimers and one TOG5 domain.

#### TOG5/GMPCPP model building

We started with the undecorated GMPCPP microtubule model PDB 6DPU and used UCSF ChimeraX^54^ and COOT^55^ to independently rigid body fit two ɑ- and two β-tubulin chains as well as a trimmed AphaFold2 model of our TOG5 construct into the corresponding density. To prevent side chains from wandering into neighboring tubulin density during refinement, we also rigid body fit six tubulin monomers and their associated ligands to form a perimeter around the two main dimers. We mutated the main tubulins from the porcine sequence to the bovine and performed minor local real space refinement in COOT before performing a global *real_space_refine* at a nominal resolution of 2.9 Å in PHENIX. Perimeter tubulin monomers were then removed from the model and *validation_cryoem* was used to calculate refinement statistics for the two-tubulin one TOG5 final model (Table 1). Molecule and map depictions presented in this work were generated using UCSF ChimeraX unless otherwise stated^54^.

**Table 1:**
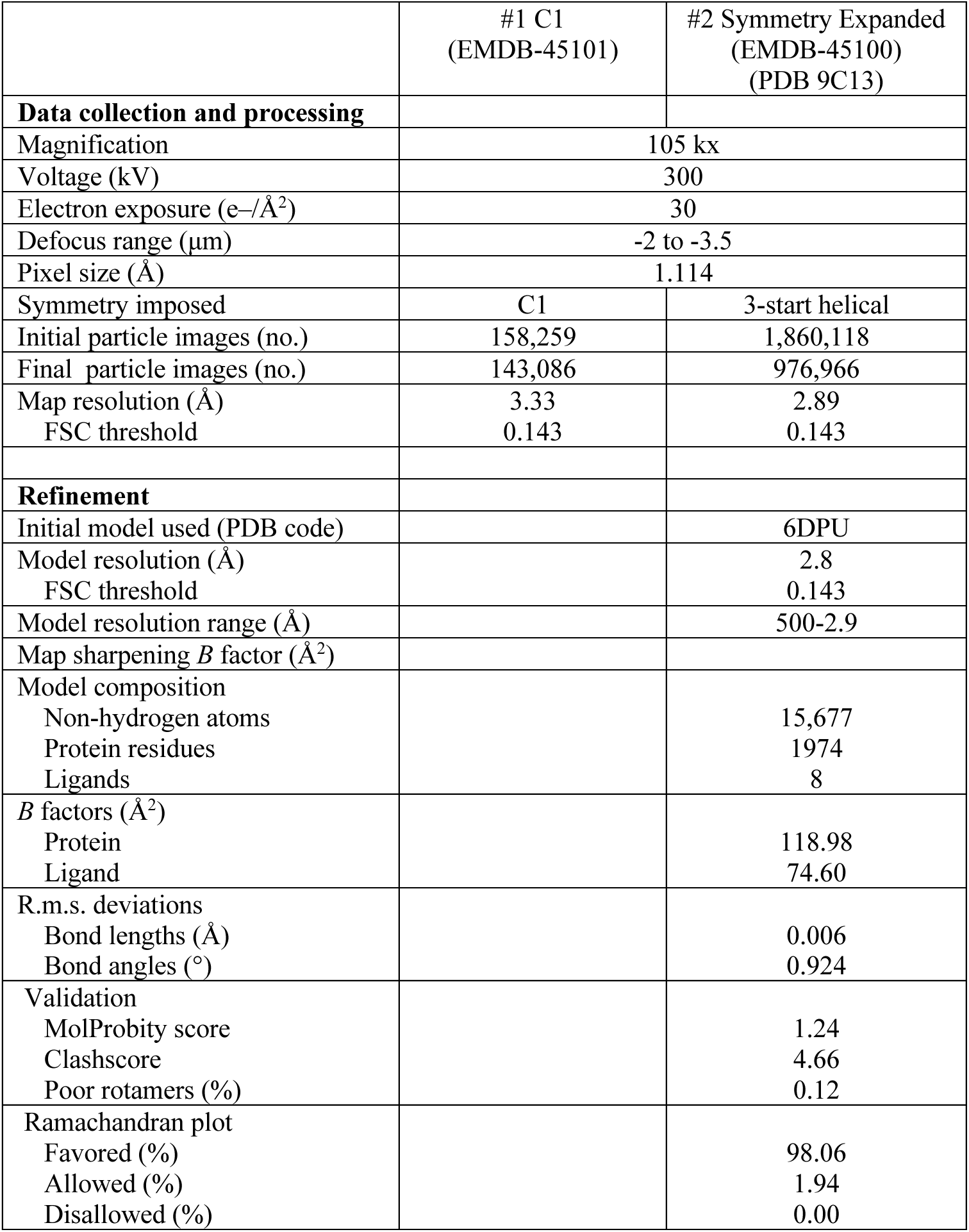
Cryo-EM data collection, refinement and validation statistics.

## Author Contributions

Collin T. McManus and Sabine Petry conceived of the project. Sabine Petry supervised the project and acquired funding. Collin T. McManus performed and analyzed all biochemical and microscopy experiments. Collin T. McManus performed crystallization screens with advice from Sophie M. Travis and Philip D. Jeffrey. Philip D. Jeffrey collected and processed crystallography data. Collin T. McManus made cryoEM grids and collected data with Sophie M. Travis. Rui Zhang processed cryoEM data, and Collin T. McManus built and refined the structural model with advice from Rui Zhang, Philip D. Jeffrey, and Sophie M. Travis. Collin T. McManus wrote the manuscript. All authors edited the manuscript.

## Acknowledgements

We would like to thank Michael Rale, Abhishek Biswas, Venecia Valdez, and Matthew Black, for their advice and assistance in generating and processing light microscopy data. We also thank Jonathan St. Ange for his help in generating reagents used in this study, and all current and former Petry Lab members for training, discussion, and reagents related to this work. C.T.M. was supported by NIH Training Grant (T32GM007388). S.M.T is supported by National Institutes of Health grants F32GM142149-01A1. R.Z. is supported by NIGMS grant 1R01GM138854. S.P. is supported by NIGMS grant 1R01GM141100-01A1.

The authors acknowledge the use of Princeton’s Imaging and Analysis Center (IAC), which is partially supported by the Princeton Center for Complex Materials (PCCM), a National Science Foundation (NSF) Materials Research Science and Engineering Center (MRSEC; DMR-2011750). This research used the AMX Beamline of the National Synchrotron Light Source II, a U.S. Department of Energy (DOE) Office of Science User Facility operated for the DOE Office of Science by Brookhaven National Laboratory under Contract No. DE-SC0012704. The Center for BioMolecular Structure (CBMS) is primarily supported by the National Institutes of Health, National Institute of General Medical Sciences (NIGMS) through a Center Core P30 Grant (P30GM133893), and by the DOE Office of Biological and Environmental Research (KP1607011). Molecular graphics and analyses performed with UCSF ChimeraX, developed by the Resource for Biocomputing, Visualization, and Informatics at the University of California, San Francisco, with support from National Institutes of Health R01-GM129325 and the Office of Cyber Infrastructure and Computational Biology, National Institute of Allergy and Infectious Diseases.

---

**S1.**
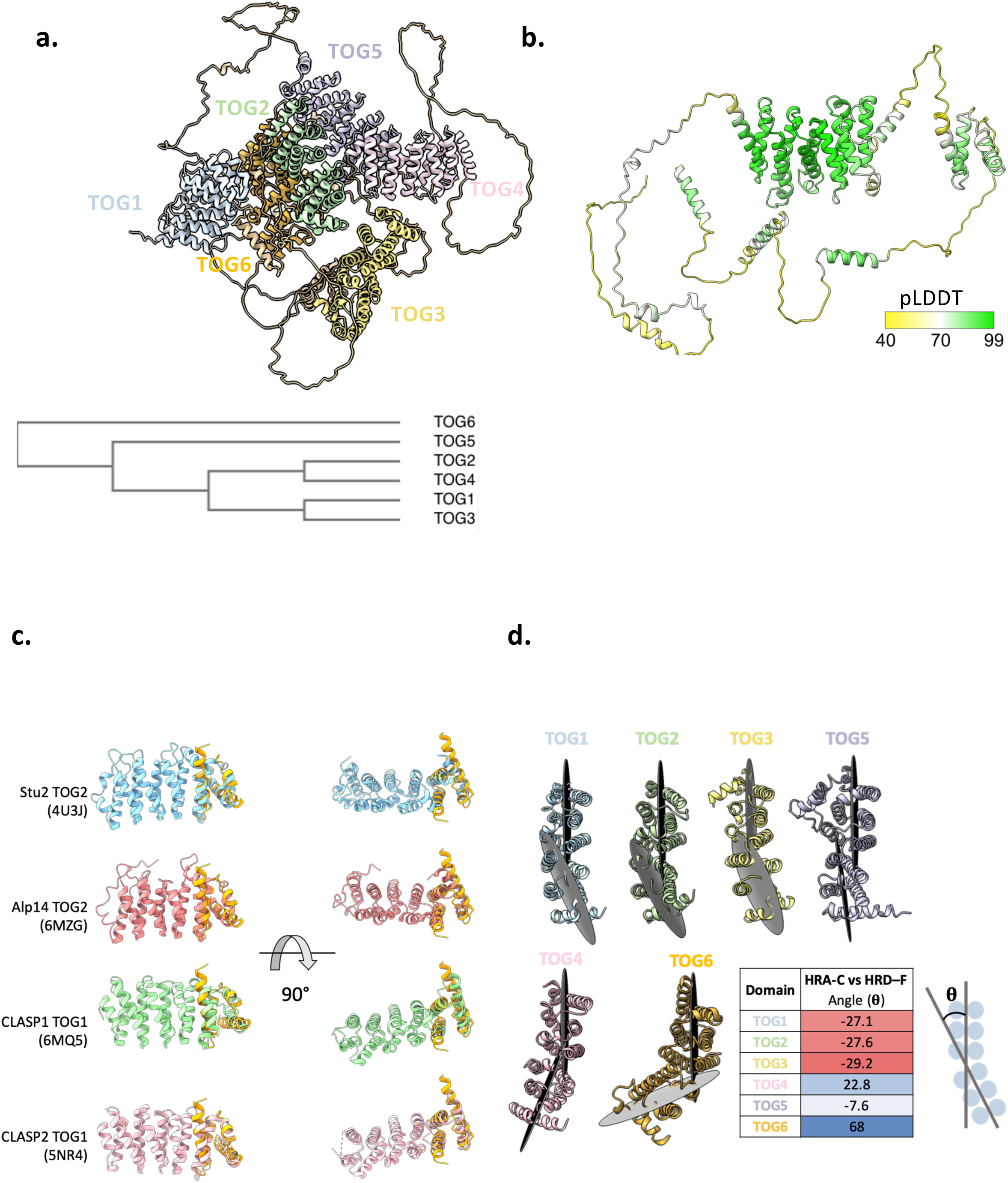
Variation in XMAP215’s TOG domains. **a)** AlphaFold2 prediction of full length XMAP215 and phylogenetic tree generated by Clustal Omega alignment of XMAP215’s six TOG domains^56^. **b)** Alphafold2 prediction of XMAP215 C-terminal region colored by pLLDT. **c)** The coordinates for atoms in the ext domain four helix bundle were used to search the PDB for structural similarity using the DALI protein structure comparison server. Among the hits were TOG domains, some of which are shown above aligned to the ext domain. N-termini are on the left, C-termini on the right. Ext domain is colored in orange. **d)** XMAP215’s TOG domains with the N-termini towards the top of the page and C-termini towards the bottom. Gray discs illustrate planes generated using heat repeats A-C and D-F for each TOG domain. Table displays measurements of angles between the two heat repeats A-C and D-F planes from B color-coded from red (negative) to blue (positive). To the right is a schematic of the angle reported, where in the case illustrated theta would be negative.

**S2.**
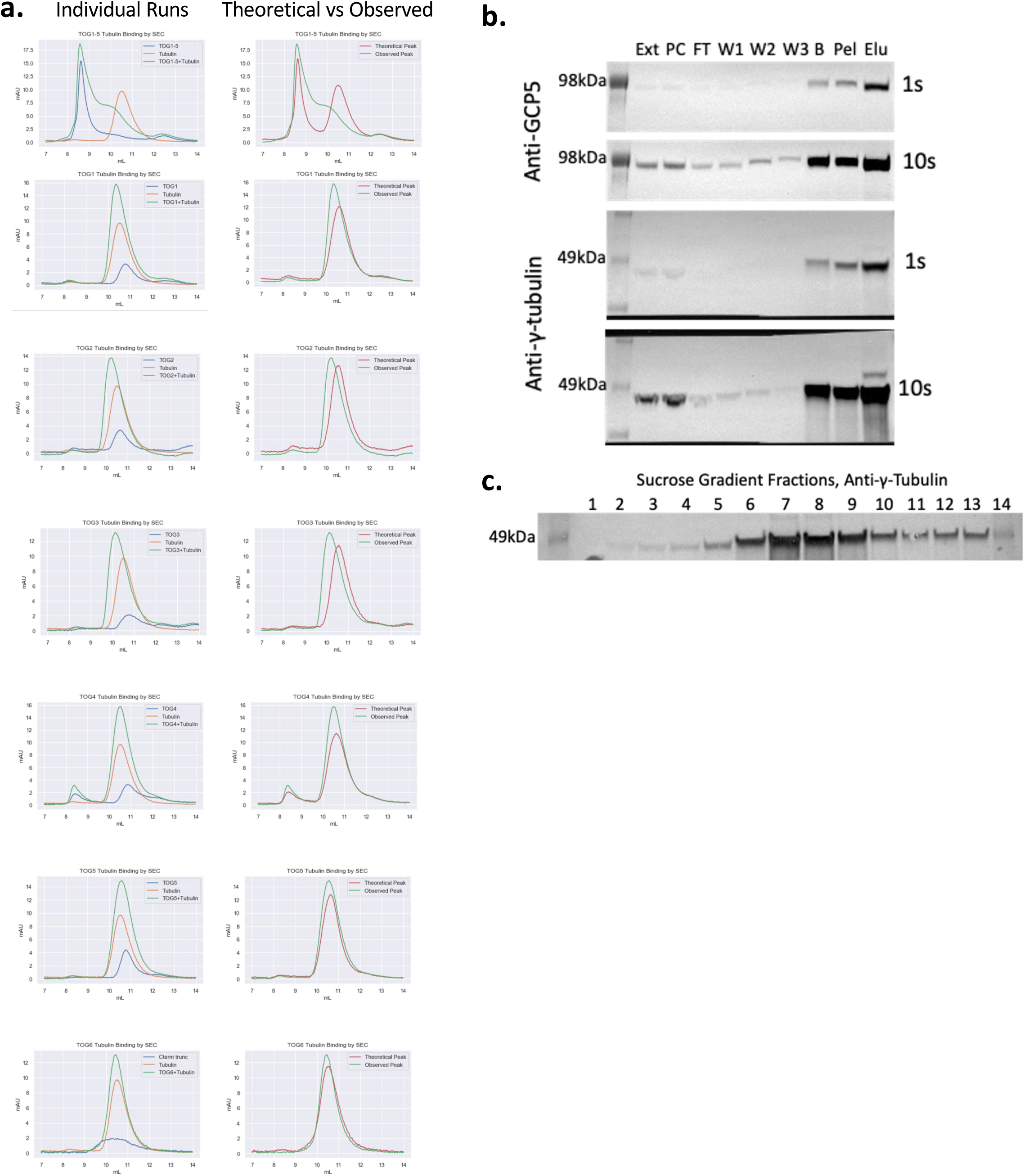
TOG6 does not bind ɑβ-tubulin. **a)** Chromatograms from gel filtration runs of GFP-tagged XMAP215 TOG1-5 and individual TOG domain constructs, tubulin, and their binding reactions. The left column shows superimposed experimental and control runs. The right column shows experimental runs superimposed with respective theoretical peaks derived by adding the traces from the control runs together. **b)** Western blot analysis of samples taken during purification of *X. laevis* endogenous γ-TuRC from oocyte extract. Ext is raw extract, PC is precleared extract, FT is extract flowthrough after incubation with γ-TuNA beads, W is wash, B is a sample taken from the γ-TuNA beads after elution of the γ-TuRC with prescission protease, Pel is a sample of a pellet of precipitate which forms during γ-TuRC concentration, and Elu is the concentrated γ-TuRC sample having been eluted from γ-TuNA beads (see methods). Exposures of 1 and 10 seconds were performed. **c)** Western blot analysis of 10-50% sucrose gradient fractionation of the elution sample from B. The peak here occurs in fractions 7 and 8.

**S3.**
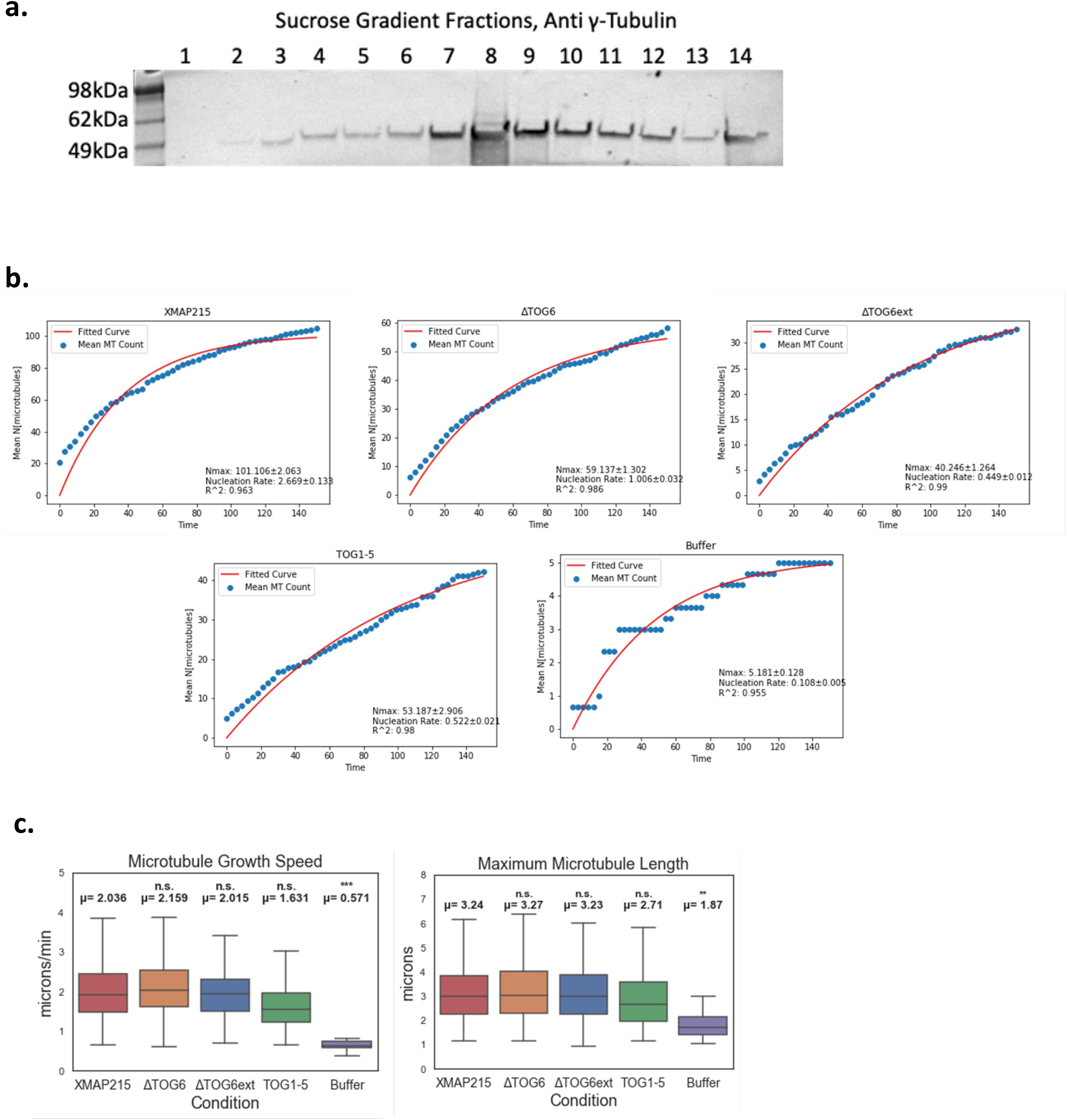
TOG6 and the ext domain do not affect microtubule polymerization. **a)** Western blot of fractions from a 10-50% sucrose gradient of biotinylated γ-TuRC using anti γ-tubulin antibody. The peak at fraction 8 was aliquoted and frozen for use in single filament nucleation reactions. **b)** Cumulative γ-TuRC nucleated microtubules over time in single filament nucleation assays, averaged across at least three reactions, for each XMAP215-GFP construct from Fig. 3b. Fitted curves are shown in red and used to derive Nmax (maximum number of microtubules nucleated) and k (nucleation rate) (see methods). R-squared represents the quality of fit of the fitted curves to the data. **c)** Box and whisker plots of microtubule growth speed and maximum length for the reactions in Fig. 3b. Plots represent data from at least three reactions. Student’s t-test was performed to compare the mean values calculated from each individual reaction for each condition. μ represents the overall average. n.s. represents p≥0.05, * represents 0.05>p>0.01, ** represents 0.01>p>0.001, and *** represents 0.001>p.

**S4.**
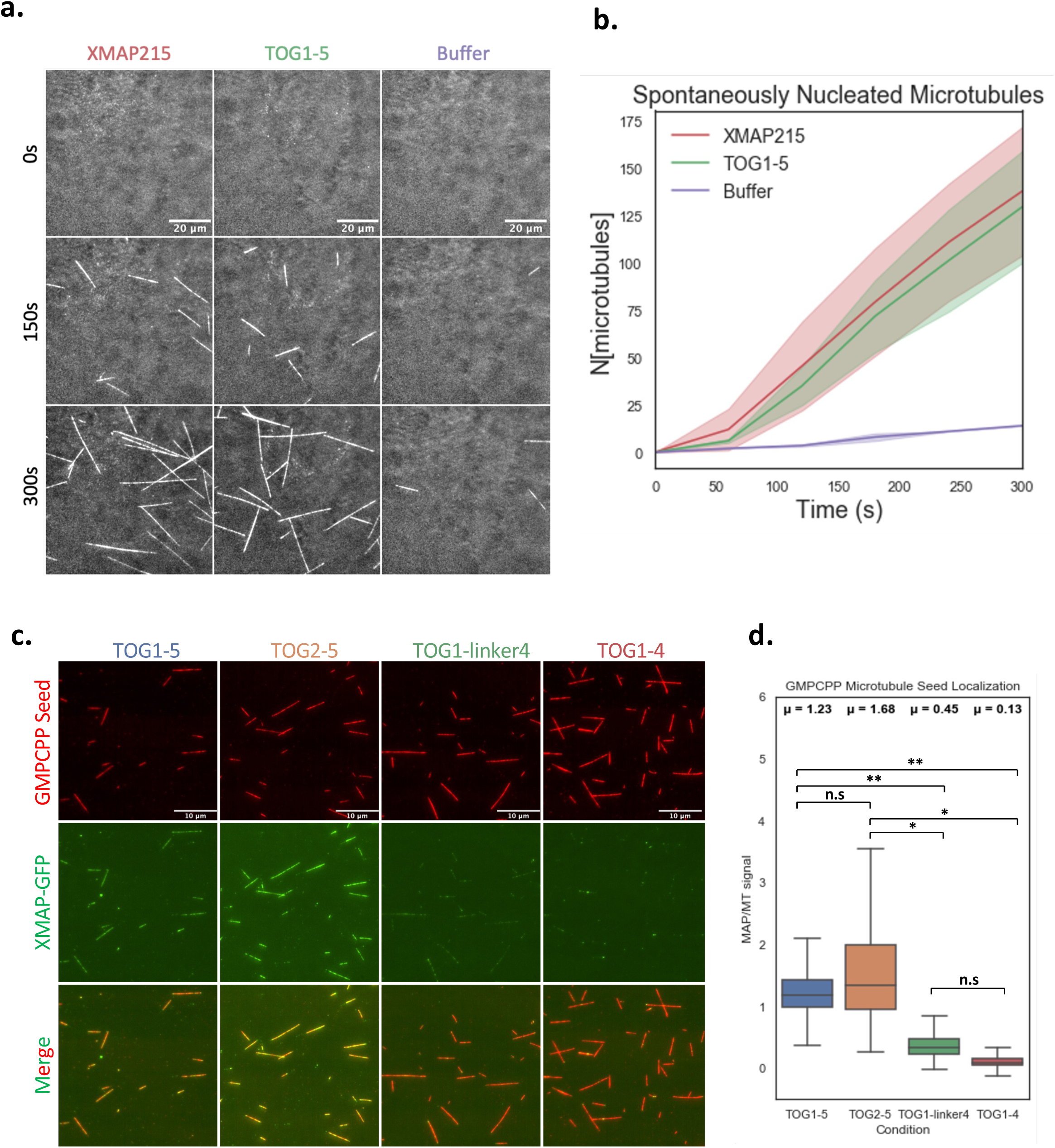
TOG1-5 is sufficient for XMAP215’s promotion of nucleation in the absence of γ-TuRC, and TOG5 helps it to bind microtubules. **a)** Time-lapse TIRF microscopy images of microtubules nucleated spontaneously using 15μM tubulin in the presence of 20nM XMAP215-GFP, TOG1-5-GFP, or control buffer. Microtubules are white against a dark background, to which a pseudo flat field correction was performed using the BioVoxxel ImageJ plugin for purposes of display ^57^. Scale bars are 20μm. **b)** Plot of microtubules present in the field view in the reactions from A over time. Lines represent averages of at least two reactions and shaded regions represent the SEM. **c)** TIRF microscopy images of Alexa568 and biotin labeled GMPCPP microtubule seeds tethered to passivated biotin-PEG coverslips by neutravidin. 200nM XMAP215-GFP constructs were allowed to incubate with the seeds for 10 minutes prior to imaging. Scale bars are 10μm. **d)** Box and whisker plots of GFP intensity (MAP) divided by tubulin (MT) intensity on segmented microtubules for each construct tested in C. Three reactions were performed for each construct. For the statistical comparisons shown, Student’s t-tests were performed using the mean MAP/MT values from each constituent reaction for each construct. n.s. represents p≥0.05, * represents 0.05>p>0.01, ** represents 0.01>p>0.001, and *** represents 0.001>p.

**S5.**
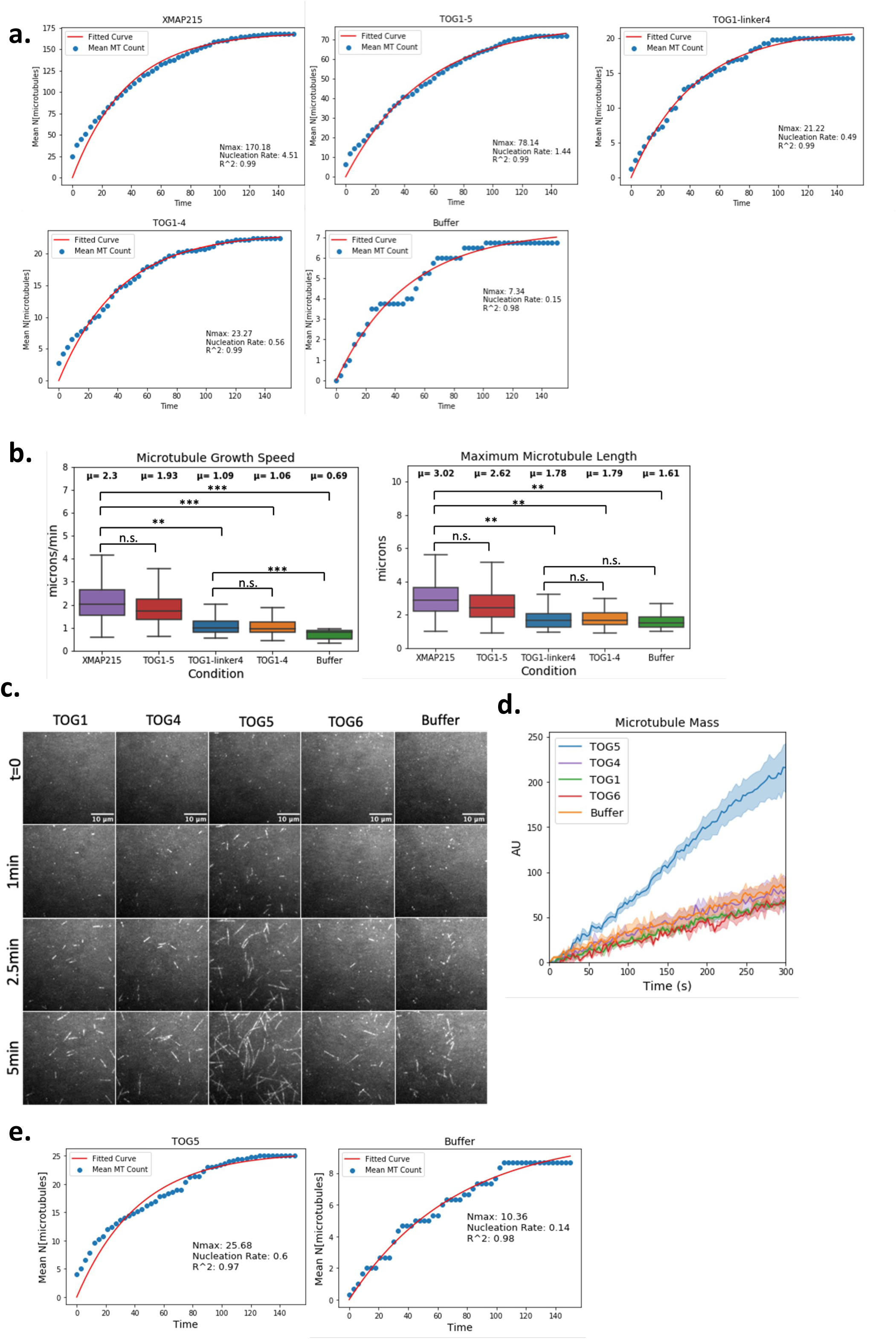
Uniquely, TOG5 is sufficient to stimulate nucleation from γ-TuRC. **a)** Cumulative γ-TuRC nucleated microtubules over time in single filament nucleation assays, averaged across at least three reactions, for each XMAP215-GFP construct from Fig. 4a. Fitted curves are shown in red and used to derive Nmax (maximum number of microtubules nucleated) and k (nucleation rate) (see methods). R-squared represents the quality of fit of the fitted curves to the data. **b)** Box and whisker plots of microtubule growth speed and maximum length for the reactions in Fig. 4a. Plots represent data from at least three reactions. Student’s t-test was performed to compare the full length XMAP215-GFP construct to each subsequent condition using mean values calculated from each individual reaction for each condition μ represents the overall average. n.s. represents p≥0.05, * represents 0.05>p>0.01, ** represents 0.01>p>0.001, and *** represents 0.001>p. **c)** Time-lapse images from single filament microtubule nucleation reactions using 7μM tubulin and 20nM TOG-GFP constructs. Microtubules are white against a dark background. **d)** Plot of microtubule intensity over time for the reactions in C. The averages of at least three reactions are plotted with the shaded region representing the corresponding SEM. **e)** Similar plots to those described in A, made for the reactions shown in C.

**S6.**
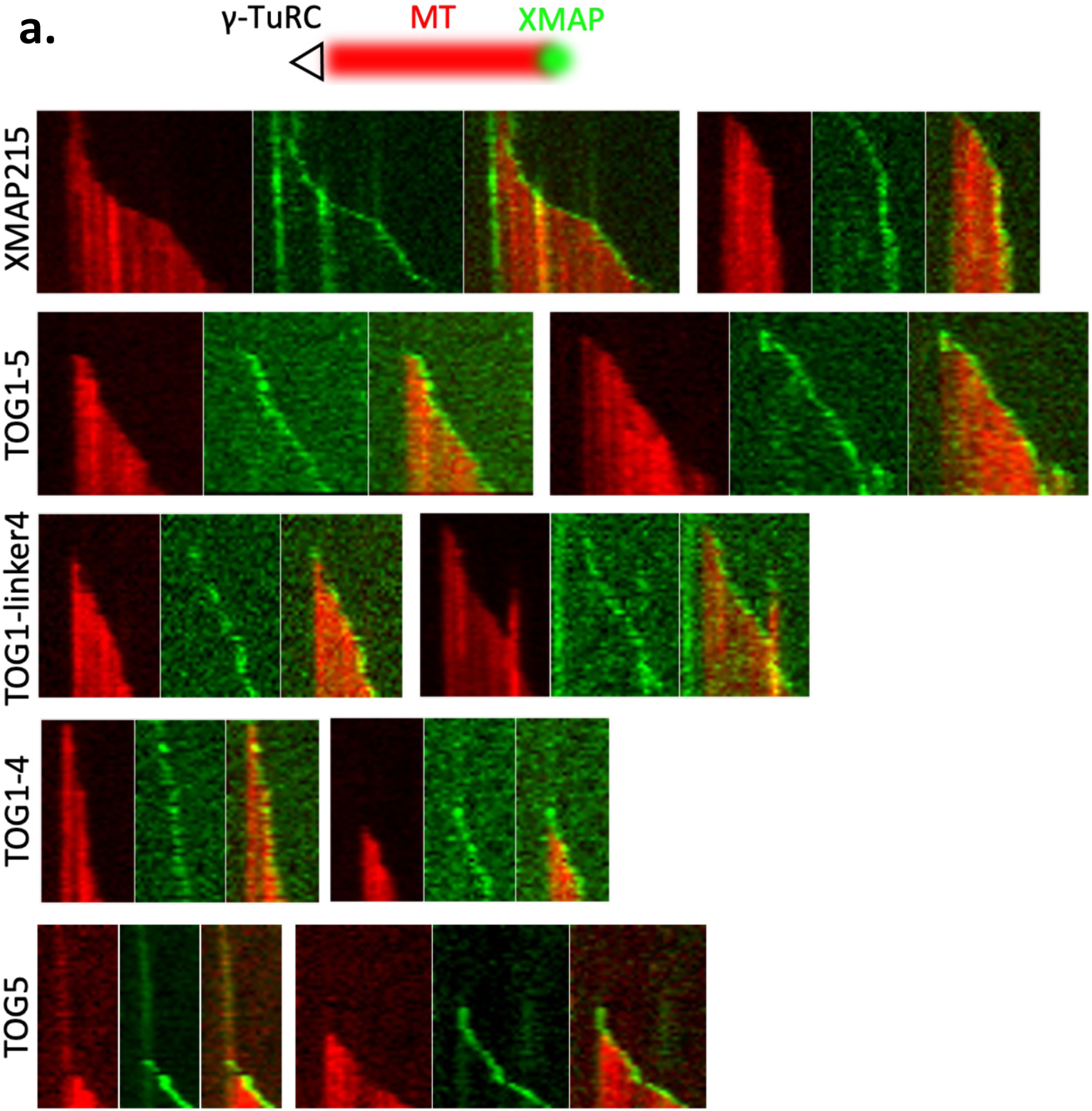
TOG5 is sufficient to track the microtubule plus end but not necessary. **a)** Representative kymographs from γ-TuRC nucleation assays shown in Fig. 4a and S5c. Microtubules are red and XMAP215-GFP constructs are green. Plus end tracking was observed consistently for XMAP215-GFP constructs containing multiple TOG domains, but was less common for TOG5-GFP.

**S7.**
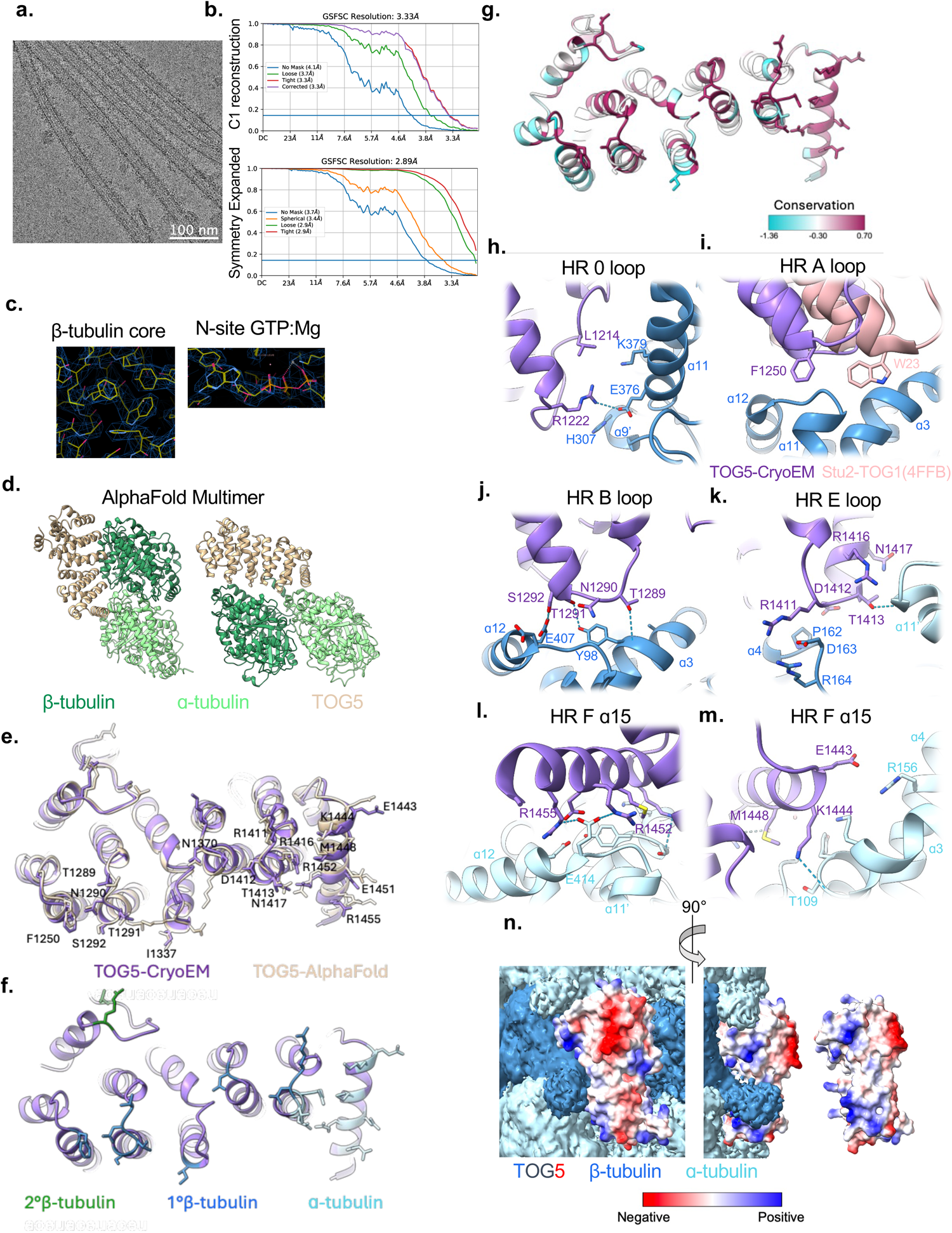
Model building and TOG5/tubulin interactions. **a)** Example micrograph of TOG5 decorated GMPCPP microtubule sample **b)** GFSC plots from local refinements of the C1 whole microtubule reconstruction and the symmetry expanded TOG5 + two tubulin masked reconstruction. **c)** Sample snapshots taken from TOG5/GMPCPP tubulin building in Coot. Bulky side chains are well-resolved, as are the GTP and Mg^2+^ ion. **d)** Two views of an AlphaFold2 multimer prediction of a TOG5/⍺β-tubulin complex. TOG5 is in tan, ⍺-tubulin in light green, and β-tubulin in dark green. **e)** TOG5 from our cryoEM structure in purple aligned with TOG5 from a TOG5/⍺β-tubulin AlphaFold2 multimer prediction in tan. Residues found to be within 5Å of tubulin residues are displayed, and those in common between the two models are labeled. **f)** Common residues from B colored according to which tubulin subunit they might interact with, dark blue for β-tubulin and light blue for ɑ-tubulin. Secondary β-tubulin interacting residues are shown in green but are unique to our cryoEM model, since AlphaFold2 assembled multiple tubulin dimers longitudinally and not laterally. **g)** Residues from C colored according to conservation, alignment in S8B. Scale is presented in terms of an entropy measure generated by AL2CO which is used by default in UCSF ChimeraX. **h-m)** Interactions of residues from C and D with tubulin labeled by structural features of TOG5. TOG5 is shown in purple, β-tubulin in dark blue, and ɑ-tubulin in light blue. The HRA panel also shows Stu2 TOG1 from Fig. 1e-f in pink. Key residues are labeled, as are ɑ-helices from tubulin subunits, and colored according to subunit. H-bonds are indicated by dashed blue lines. Heteroatoms are colored red for oxygen, dark blue for nitrogen, and yellow for sulfur. **n)** The C1 reconstruction from Fig. 5a colored according to tubulin subunit. TOG5 density is not shown and replaced with a surface model colored according to charge. Density consistent with tubulin tails can be seen protruding from the lattice and are also colored according to subunit. The rightmost image shows a sideview of the TOG5 surface model with the tubulin density removed.

**S8.**
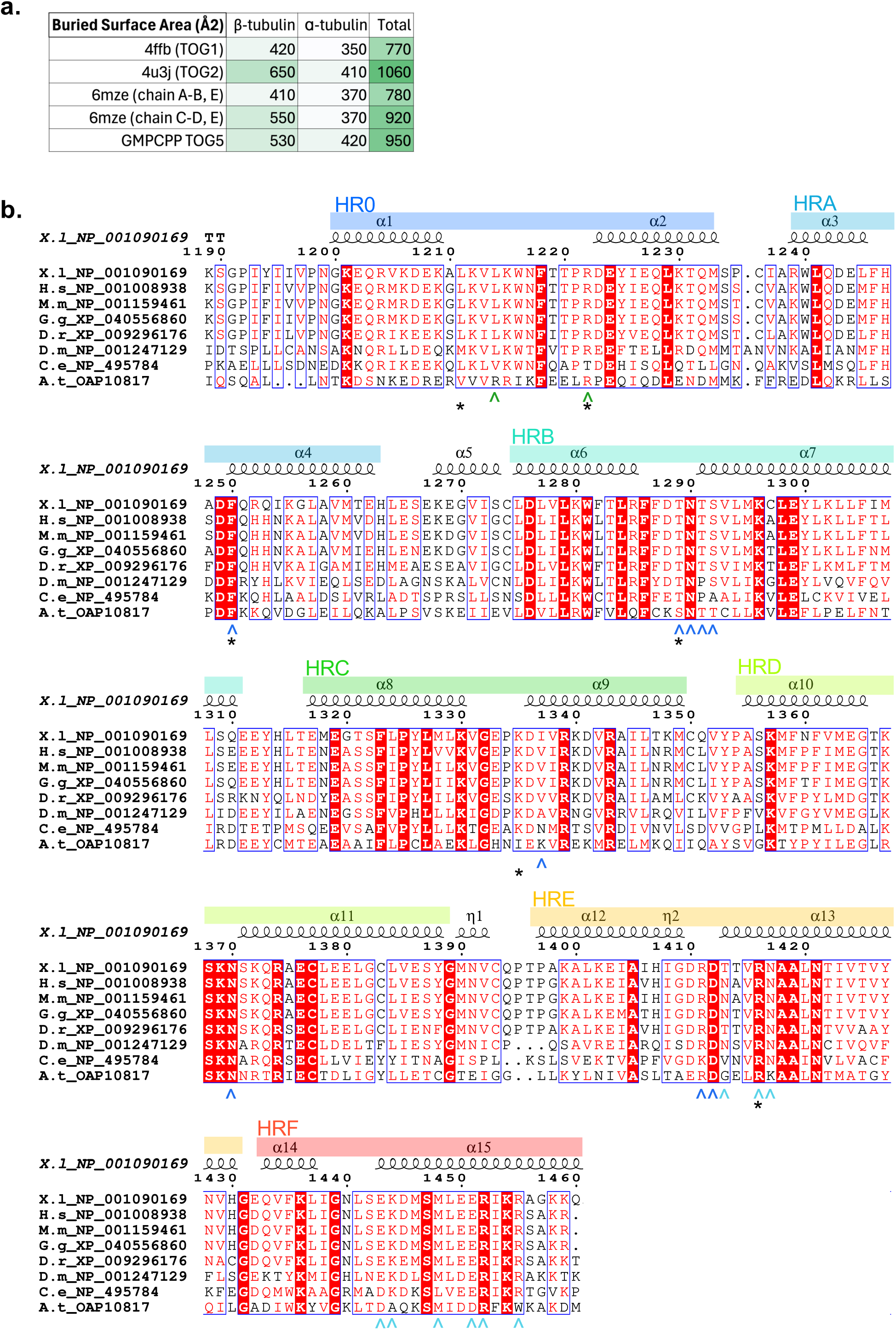
TOG5 binds straight tubulin analogously to TOGs 1 and 2 binding bent tubulin, and the residues involved are conserved. **a)** A table showing calculations made in UCSF ChimeraX for buried surface area between TOG domains and tubulin subunits in the structures named according to PDB accession, and the GMPCPP tubulin/TOG5 model presented in this work. Buried area is colored from white to green according to magnitude. **b)** Multiple sequence alignment of TOG5 from common model organisms labeled by initials and protein accession number. Organisms are *Xenopus laevis, Homo sapiens, Mus musculus, Gallus gallus, Danio rerio, Drosophila melanogaster, Caenorhabditis elegans,* and *Arabidopsis thalania.* Numbering is according to XMAP215 from *Xenopus laevis*. Secondary structure elements are shown at the top and highlighted with colored bars according to HEAT repeat, depicted in Fig. 5b. Carrots below the alignment correspond with residues displayed and colored in S7D. Asterisks (*) indicate residues corresponding to or near those identified in S7D-E which have been previously mutated in TOG domains and found to be important for tubulin binding (Ayaz et al., 2012, Bynes et al., 2017). The alignment was generated using the CLUSTAL implementation in SnapGene and displayed using ESPript - https://espript.ibcp.fr.

**Movie S1.** Live assays as depicted as stills in Fig. 3b.

**Movie S2.** Live assays as depicted as stills in Fig. 4a.

**Movie S3.** Live assays as depicted as stills in Fig. S4a.

**Movie S4.** Live assays as depicted as stills in Fig. S5c.

## References

1. Petry, S. & Vale, R. D. Microtubule nucleation at the centrosome and beyond. Nat. Cell Biol. 17, 1089–1093 (2015).

2. Wu, J. & Akhmanova, A. Microtubule-Organizing Centers. (2017).

3. Travis, S. M., Mahon, B. P. & Petry, S. How Microtubules Build the Spindle Branch by Branch. Annu. Rev. Cell Dev. Biol. 38, 1–23 (2022).

4. Thawani, A., Kadzik, R. S. & Petry, S. XMAP215 is a microtubule nucleation factor that functions synergistically with the γ-tubulin ring complex. Nat. Cell Biol. 20, 575–585 (2018).

5. Ali, A., Vineethakumari, C., Lacasa, C. & Lüders, J. Microtubule nucleation and γTuRC centrosome localization in interphase cells require ch-TOG. Nat. Commun. 14, 289 (2023).

6. Zhu, Z. et al. Multifaceted modes of γ-tubulin complex recruitment and microtubule nucleation at mitotic centrosomes. J. Cell Biol. 222, e202212043 (2023).

7. Kollman, J. M., Polka, J. K., Zelter, A., Davis, T. N. & Agard, D. A. Microtubule nucleating γ-TuSC assembles structures with 13-fold microtubule-like symmetry. Nature 466, 879–882 (2010).

8. Kollman, J. M. et al. Ring closure activates yeast γTuRC for species-specific microtubule nucleation. Nat. Struct. Mol. Biol. 22, 132–137 (2015).

9. Wieczorek, M. et al. Asymmetric Molecular Architecture of the Human γ-Tubulin Ring Complex. Cell 180, 165–175.e16 (2020).

10. Consolati, T. et al. Microtubule Nucleation Properties of Single Human γTuRCs Explained by Their Cryo-EM Structure. Dev. Cell 53, 603–617.e8 (2020).

11. Liu, P. et al. Insights into the assembly and activation of the microtubule nucleator γ-TuRC. Nature 578, 467–471 (2020).

12. Thawani, A. et al. The transition state and regulation of γ-TuRC-mediated microtubule nucleation revealed by single molecule microscopy. eLife 9, e54253 (2020).

13. Rale, M. J., Romer, B., Mahon, B. P., Travis, S. M. & Petry, S. The conserved centrosomin motif, γTuNA, forms a dimer that directly activates microtubule nucleation by the γ-tubulin ring complex (γTuRC). eLife 11, e80053 (2022).

14. Aher, A., Urnavicius, L., Xue, A., Neselu, K. & Kapoor, T. M. Structure of the γ-tubulin ring complex-capped microtubule.

15. Brito, C., Serna, M., Guerra, P., Llorca, O. & Surrey, T. Transition of human γ-tubulin ring complex into a closed conformation during microtubule nucleation. Science 383, 870–876 (2024).

16. Dendooven, T. et al. Structure of the native γ-tubulin ring complex capping spindle microtubules. Nat. Struct. Mol. Biol. 1–11 (2024) doi:10.1038/s41594-024-01281-y.

17. Popov, A. V. XMAP215 regulates microtubule dynamics through two distinct domains. EMBO J. 20, 397–410 (2001).

18. Popov, A. V., Severin, F. & Karsenti, E. XMAP215 Is Required for the Microtubule-Nucleating Activity of Centrosomes. Curr. Biol. 12, 1326–1330 (2002).

19. Brouhard, G. J. et al. XMAP215 Is a Processive Microtubule Polymerase. Cell 132, 79–88 (2008).

20. Al-Bassam, J. et al. Fission yeast Alp14 is a dose-dependent plus end–tracking microtubule polymerase. Mol. Biol. Cell 23, 2878–2890 (2012).

21. Al-Bassam, J. & Chang, F. Regulation of microtubule dynamics by TOG-domain proteins XMAP215/Dis1 and CLASP. Trends Cell Biol. 21, 604–614 (2011).

22. Slep, K. C. & Vale, R. D. Structural Basis of Microtubule Plus End Tracking by XMAP215, CLIP-170, and EB1. Mol. Cell 27, 976–991 (2007).

23. Ayaz, P., Ye, X., Huddleston, P., Brautigam, C. A. & Rice, L. M. A TOG:αβ-tubulin Complex Structure Reveals Conformation-Based Mechanisms for a Microtubule Polymerase. Science 337, 857–860 (2012).

24. Ayaz, P. et al. A tethered delivery mechanism explains the catalytic action of a microtubule polymerase. eLife 3, e03069 (2014).

25. Al-Bassam, J., Larsen, N. A., Hyman, A. A. & Harrison, S. C. Crystal Structure of a TOG Domain: Conserved Features of XMAP215/Dis1-Family TOG Domains and Implications for Tubulin Binding. Structure 15, 355–362 (2007).

26. Byrnes, A. E. & Slep, K. C. TOG–tubulin binding specificity promotes microtubule dynamics and mitotic spindle formation. J. Cell Biol. 216, 1641–1657 (2017).

27. Fox, J. C., Howard, A. E., Currie, J. D., Rogers, S. L. & Slep, K. C. The XMAP215 family drives microtubule polymerization using a structurally diverse TOG array. Mol. Biol. Cell 25, 2375–2392 (2014).

28. Howard, A. E., Fox, J. C. & Slep, K. C. Drosophila melanogaster Mini Spindles TOG3 Utilizes Unique Structural Elements to Promote Domain Stability and Maintain a TOG1- and TOG2-like Tubulin-binding Surface. J. Biol. Chem. 290, 10149–10162 (2015).

29. Nithianantham, S. et al. Structural basis of tubulin recruitment and assembly by microtubule polymerases with tumor overexpressed gene (TOG) domain arrays. eLife 7, e38922 (2018).

30. Widlund, P. O., et al. XMAP215 polymerase activity is built by combining multiple tubulin-binding TOG domains and a basic lattice-binding region. Proc. Natl. Acad. Sci. 108, 2741–2746 (2011).

31. Slep, K. C. A Cytoskeletal Symphony: Owed to TOG. Dev. Cell 46, 5–7 (2018).

32. King, B. R. et al. XMAP215 and γ-tubulin additively promote microtubule nucleation in purified solutions. Mol. Biol. Cell 31, 2187–2194 (2020).

33. Gunzelmann, J. et al. The microtubule polymerase Stu2 promotes oligomerization of the γ-TuSC for cytoplasmic microtubule nucleation. eLife 7, e39932 (2018).

34. Flor-Parra, I., Iglesias-Romero, A. B. & Chang, F. The XMAP215 Ortholog Alp14 Promotes Microtubule Nucleation in Fission Yeast. Curr. Biol. 28, 1681–1691.e4 (2018).

35. Roostalu, J., Cade, N. I. & Surrey, T. Complementary activities of TPX2 and chTOG constitute an efficient importin-regulated microtubule nucleation module. Nat. Cell Biol. 17, 1422–1434 (2015).

36. Hood, F. E. et al. Coordination of adjacent domains mediates TACC3–ch-TOG–clathrin assembly and mitotic spindle binding. J. Cell Biol. 202, 463–478 (2013).

37. Rostkova, E., Burgess, S. G., Bayliss, R. & Pfuhl, M. Solution NMR assignment of the C-terminal domain of human chTOG. Biomol. NMR Assign. 12, 221–224 (2018).

38. Jumper, J. et al. Highly accurate protein structure prediction with AlphaFold. Nature 596, 583–589 (2021).

39. Sandoval, I. V., MacDonald, E., Jameson, J. L. & Cuatrecasas, P. Role of nucleotides in tubulin polymerization: effect of guanylyl 5’-methylenediphosphonate. Proc. Natl. Acad. Sci. U. S. A. 74, 4881–4885 (1977).

40. Geyer, E. A., Miller, M. P., Brautigam, C. A., Biggins, S. & Rice, L. M. Design principles of a microtubule polymerase. eLife 7, e34574 (2018).

41. Cook, B. D., Chang, F., Flor-Parra, I. & Al-Bassam, J. Microtubule polymerase and processive plus-end tracking functions originate from distinct features within TOG domain arrays. Mol. Biol. Cell 30, 1490–1504 (2019).

42. Zhang, R. & Nogales, E. A new protocol to accurately determine microtubule lattice seam location. J. Struct. Biol. 192, 245–254 (2015).

43. Valdez, V., Ma, M., Gouveia, B., Zhang, R. & Petry, S. HURP facilitates spindle assembly by stabilizing microtubules and working synergistically with TPX2. 2023.12.18.571906 Preprint at 10.1101/2023.12.18.571906 (2023).

44. Brouhard, G. J. & Rice, L. M. Microtubule dynamics: an interplay of biochemistry and mechanics. Nat. Rev. Mol. Cell Biol. 19, 451–463 (2018).

45. Gard, D. L., Becker, B. E. & Josh Romney, S. MAPping the Eukaryotic Tree of Life: Structure, Function, and Evolution of the MAP215⧸Dis1 Family of Microtubule-Associated Proteins. in International Review of Cytology vol. 239 179–272 (Elsevier, 2004).

46. Romer, B. et al. Conformational states of the microtubule nucleator, the γ-tubulin ring complex. 2023.12.19.572162 Preprint at 10.1101/2023.12.19.572162 (2023).

47. Wieczorek, M., Huang, T.-L., Urnavicius, L., Hsia, K.-C. & Kapoor, T. M. MZT Proteins Form Multi-Faceted Structural Modules in the γ-Tubulin Ring Complex. Cell Rep. 31, 107791 (2020).

48. Sabo, J. et al. CKAP5 enables formation of persistent actin bundles templated by dynamically instable microtubules. Curr. Biol. 34, 260–272.e7 (2024).

49. Wieczorek, M., Bechstedt, S., Chaaban, S. & Brouhard, G. J. Microtubule-associated proteins control the kinetics of microtubule nucleation. Nat. Cell Biol. 17, 907–916 (2015).

50. Schindelin, J., et al. Fiji: an open-source platform for biological-image analysis. Nat. Methods 9, 676–682 (2012).

51. Thevenaz, P., Ruttimann, U. E. & Unser, M. A pyramid approach to subpixel registration based on intensity. IEEE Trans. Image Process. 7, 27–41 (1998).

52. Sharma, V. ImageJ plugin HyperStackReg V5.6. Zenodo 10.5281/zenodo.2252521 (2018).

53. Punjani, A., Rubinstein, J. L., Fleet, D. J. & Brubaker, M. A. cryoSPARC: algorithms for rapid unsupervised cryo-EM structure determination. Nat. Methods 14, 290–296 (2017).

54. Meng, E. C. et al. UCSF ChimeraX: Tools for structure building and analysis. Protein Sci. 32, e4792 (2023).

55. Emsley, P., Lohkamp, B., Scott, W. G. & Cowtan, K. Features and development of *Coot*. Acta Crystallogr. D Biol. Crystallogr. 66, 486–501 (2010).

56. Sievers, F. & Higgins, D. G. Clustal Omega for making accurate alignments of many protein sequences. Protein Sci. 27, 135–145 (2018).

57. Brocher, J. biovoxxel/BioVoxxel-Toolbox. (2024).

